# Neural substrates of body ownership and agency during voluntary movement

**DOI:** 10.1101/2022.07.23.501221

**Authors:** Z Abdulkarim, A Guterstam, Z Hayatou, HH Ehrsson

**Affiliations:** Department of Neuroscience, Karolinska Institutet, Stockholm, Sweden; Department of Clinical Neuroscience, Karolinska Institutet, Stockholm, Sweden; Université Paris-Saclay, CNRS, Institut Des Neurosciences Paris-Saclay, 91190 Gif-sur-Yvette, France

## Abstract

Body ownership and the sense of agency are two central aspects of bodily self-consciousness. While multiple neuroimaging studies have investigated the neural correlates of body ownership and agency in isolation, few have investigated their relationship during voluntary movement when such experiences naturally combine. By eliciting the moving rubber hand illusion with active or passive finger movements during functional magnetic resonance imaging, we isolated activations reflecting the sense of body ownership and agency, respectively, as well as their interaction, and assessed their overlap and anatomical segregation. We found that perceived hand ownership was associated with activity in premotor, posterior parietal and cerebellar regions whereas the sense of agency over the hand’s movements was related to activity in the dorsal premotor cortex and superior temporal cortex. Moreover, one section of the dorsal premotor cortex showed overlapping activity for ownership and agency, and somatosensory cortical activity reflected the interaction of ownership and agency with higher activity when both agency and ownership was experienced. We further found that activations previously attributed to agency in the left insular cortex and right temporoparietal junction reflected the synchrony or asynchrony of the visuo-proprioceptive stimuli rather than agency. Collectively, these results identify the neural bases of agency and ownership during voluntary movement. Although the neural representations of these two experiences are largely distinct, there are functional neuroanatomical overlap and interactions during their combination, which has bearing on theories on bodily self-consciousness.

## Introduction

When you raise your arm, you automatically experience that it was you who caused the arm to lift and that the moving arm is your own. These two experiences blend so naturally during everyday voluntary behavior that we rarely think of them as distinct. However, in philosophy, cognitive science, and cognitive neuroscience there is a long tradition to study the sense of being in control of and causing bodily action through volition – the *sense of agency* (Haggard, 2017; Jeannerod, 2003) – and the immediate perceptual experience of limbs and body parts as one’s own – the *sense of body ownership* (H. H. Ehrsson, 2020; Petkova & Ehrsson, 2010) as distinct processes. Body ownership and agency are both considered to be fundamental aspects of self-consciousness and critical for defining what it is to be a conscious embodied agent distinct from the environment. However, most previous studies have focused on these two experiences in isolation using different experimental paradigms, so little is known about how they combine during voluntary movement.

Body ownership is considered to depend on the integration of visual, somatosensory and other sensory bodily signals into coherent multisensory percepts of the own body through mechanisms of multisensory integration (H. H. Ehrsson et al., 2004; H. H. Ehrsson, 2020), whereas agency relates to the association between voluntary action and outcome and has been linked to the match between the expected sensory consequences of movement and their sensory feedback (Frith et al., 2000) and the experience of volition during voluntary movement (Haggard, 2017). Previous fMRI studies have identified brain areas associated with the sense of body ownership and sense of agency, where body ownership is associated with activity in a set of premotor-parieto-cerebellar regions (H. H. Ehrsson et al., 2004, 2005; Gentile et al., 2015; Guterstam et al., 2013; Limanowski & Blankenburg, 2016) and agency related to activations in the right inferior parietal cortex, temporoparietal junction, pre-supplementary motor area (pre-SMA), insula (Chambon et al., 2013; David et al., 2008; Farrer et al., 2003; Farrer & Frith, 2002; Schnell et al., 2007; Yomogida et al., 2010), superior temporal gyrus (STG) (Nahab et al., 2011; Uhlmann et al., 2020) and the left primary sensorimotor cortex (Sperduti et al., 2011). However, the agency studies have focused on agency over external sensory events that happens as a consequence of bodily movement rather than agency experienced directly over one’s moving limbs, and the body ownership studies have not investigated movement (but see Tsakiris et al 2010). The precise functional neuroanatomical relationship between ownership and agency during simple voluntary movement is therefore unclear.

Here we used the rubber hand illusion (RHI) (Botvinick & Cohen, 1998) elicited by finger movements – the moving RHI (Kalckert & Ehrsson, 2012) – to investigate the neural bases of body ownership and agency within a single functional magnetic resonance imaging (fMRI) paradigm. To elicit this bodily illusion, the participants perform a repetitive finger movement with their hidden index finger while they observe a rubber hand placed in full view making the corresponding finger movements. After a few synchronous movements, the participants start to experience the moving rubber hand as their own and that they are directly controlling its movements voluntarily (Kalckert & Ehrsson, 2012, 2014). By manipulating the relative timing of the real and rubber hand’s finger movements (synchrony or asynchrony), the type of movement (active or passive), and spatial-anatomical orientation of the rubber hand with respect to the and real hand (congruent or incongruent) the sense of body ownership and agency can be individually manipulated (Kalckert & Ehrsson, 2012). Thus, we implemented a 2×2×2 factorial within-subjects experimental design with these three factors to identify active neuronal populations that reflect body ownership, agency and their potential interaction. We hypothesized that ownership and agency should be associated with activity in different neural circuits in line with previous studies, but also, that their combination should be associated with overlapping and stronger activation in certain frontoparietal regions due to the integration of the two sensations.

## Materials and Methods

### Participants

Thirty healthy volunteers were recruited for the experiment. One of the participants cancelled their participation last minute, leaving 29 participants that completed the experiment (15 males, 14 females, mean age 28 ± 5). The number of participants recruited was based on previous similar studies on body illusions (Preston & Ehrsson, 2016) as well as another fMRI studies with a similar 2×2×2 factorial design and eight conditions in a block design (Kilteni & Ehrsson, 2020). All the participants were right-handed, assessed with the Edinburgh handedness inventory (Oldfield, 1971). The participants had normal or corrected to normal vision and had no history of neurological or psychiatric illness. Informed consent was obtained prior to the experiment. The experiment was conducted according to the declaration of Helsinki and was approved by the Swedish Ethical Review Authority.

### Moving rubber hand illusion setup

The moving rubber hand illusion setup in its original design is a vertical setup in which the participants real hand is placed under a small table over which the rubber hand is placed (Kilteni & Ehrsson, 2012), but the illusion also works in other spatial arrangement as long as the rubber hand is presented close to the real hand within peri-hand space (approx. within a 30-40 cm distance), and we took this into consideration when re-designing the setup for the current fMRI study. The vertical setup did not fit inside the constrained space of the modern GE 3T MR scanner we use, so a horizontal version of the moving rubber hand illusion setup had to be designed. Importantly, the setup had to be able to rapidly switch between active and passive movements of the participant’s index finger, as well as between synchronous and asynchronous movements of the participants index finger and the index finger of the rubber hand. To achieve this, a new mechanical design consisting of two levers, two supports and a plastic pin was developed (Fig. 1, Panels A-D). By removing the plastic pin connecting the levers the movements of the participants index finger could be decoupled from the movements of the index finger of the rubber hand. By having the experimenter push the lever beneath the index finger up, the fingers could be passively moved. The “rubber hand” used in our experiment was in fact a wooden hand with flexible joints (HAY design brand, 31 cm model; similar to (Kilteni & Ehrsson, 2012). All joints but the metacarpophalangeal (MCP) joint of the index finger were fixated with glue, thus only permitting movement in the MCP joint. The rubber hand was covered with a grey nitrile glove, occluding the fact that it was a wooden hand and giving it the impression of being more humanoid.

**Figure 1.**
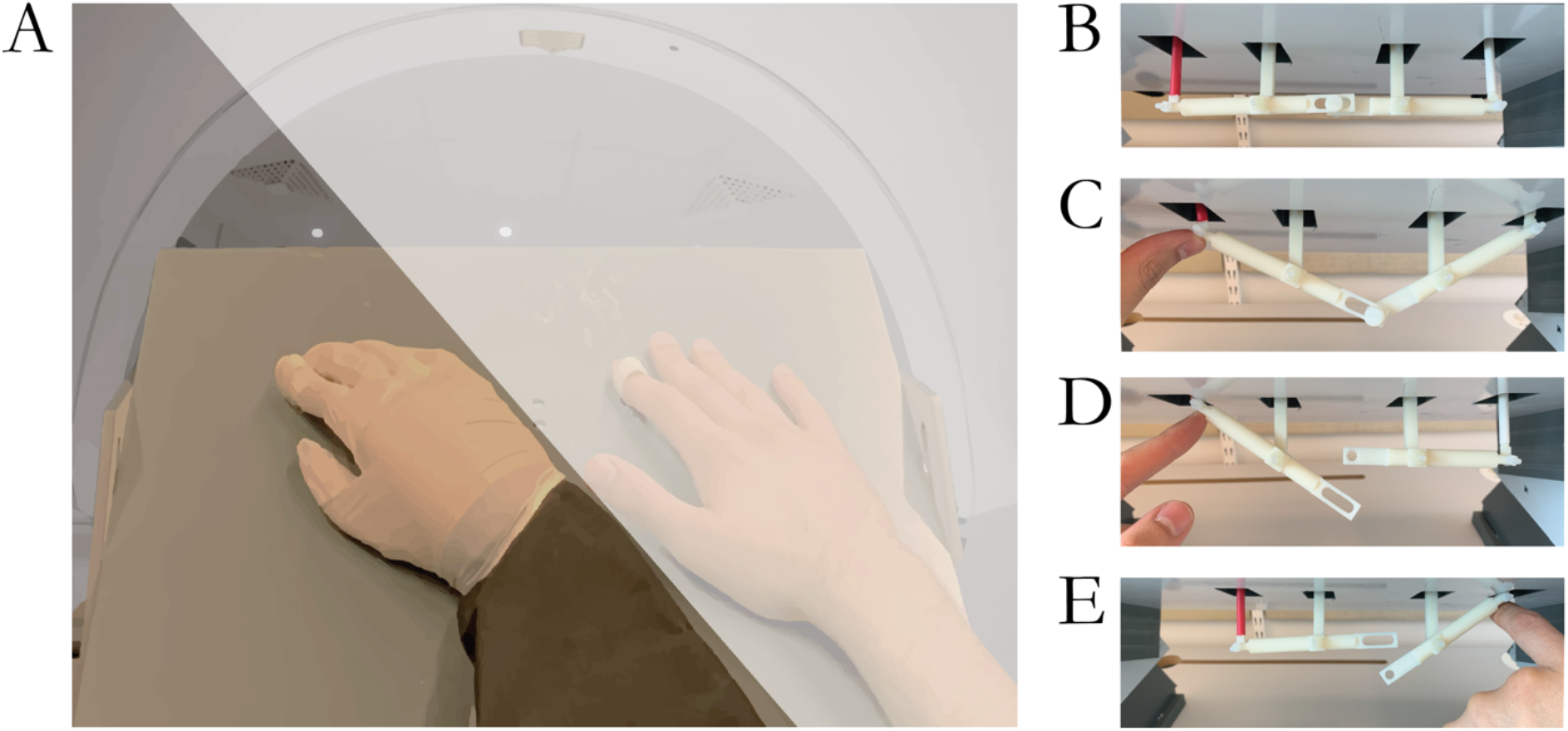
**A**. Depicts a montage of what the participants would see laying inside the MR-scanner. The white semi-opaque field illustrates the dark cloth used to cover the participant’s real right hand from view. The participants hand and the rubber hand are seen resting on a small table. The index finger of the rubber hand as well as the participant’s hand is placed inside a plastic ring, which is connected to the two most lateral vertical rods seen in panel B-E. Panel **B-E** illustrates the levers of the moving rubber hand illusion setup under the table that moved the index finger of the participant and the rubber hand. In **B**, the levers are in a relaxed position with the index finger of the rubber hand and the participant’s hand resting on the table. In **C**, both the participant’s index finger and the index finger of the rubber hand is lifted off the table. The two levers are connected to each other through a pin. In this configuration the participants could lift their index finger, which would simultaneously lift the index finger of the rubber hand (active synchronous conditions), or the experimenter could push the index finger of the participant up by pressing on the rod underneath the participants index finger (as seen in the image; passive synchronous condition). In **D & E**, the two fingers have been decoupled by removing the pin holding the two levers together. In this configuration, the index finger of the rubber hand and the participants hand could be moved independently by the experimenter; delayed movements (approx. 0.5s) of the rubber hand’s index in the asynchronous conditions (active and passive asynchronous conditions).

### Procedures

The participant lay comfortably in a supine position on the MR scanner bed wearing earplugs and headphones over the earplugs to protect the participant’s hearing from the scanner noise while allowing them to hear the instructions through the headphones. The participant’s head was tilted approximately 30 degrees using a custom-made wooden wedge under the head coil along with foam pads inside the head coil. The tilting of the head allowed the participants to see through the openings on the head coil and view their body from a natural (first-person) perspective. With the participant supine, a small custom-made table was placed over their abdomen (fixed to the scanner bed). The participant’s right hand was placed on the right side of this table, and the rubber hand was placed on the left side of this table, with the index finger of the rubber hand placed 15 cm to the left of the index finger of the participant’s real hand (Fig. 1, Panel A). The participant’s real right arm and hand was occluded by taping a dark cloth to the table and then to the roof of the MR scanner bore, thus completely hiding the participants real hand from sight (Fig. 1, Panel A). The participant’s right elbow was supported with a pillow in order to have the participants lay comfortably and not having to strain or actively maintain their arm in the required position but make it possible for them to have the arm in a relaxed posture. The rubber hand and the participant’s real hand were placed parallel to each other, with the same rotations of approximately 20 degrees counterclockwise from the participants perspective, which gave the impression of the rubber hand originating from the insertion of the participants real arm to the torso. The participants right index finger was placed inside a small plastic ring which was connected to a rod that in turn was connected to a lever below the table (Fig. 1). The index finger of the rubber hand was placed in an identical plastic ring and in turn connected via a separate rod to a second lever under the table. This setup allowed us to manipulate the synchrony of the movements between the rubber hand and the real hand by coupling (synchrony) or decoupling (i.e., removing the plastic pin connecting the two levers) the rubber hand from the participant’s hand (Fig 1B-E). This decoupling allowed the index finger of the rubber hand and the participant’s hand to move independently, and thus the experimenter could move the index finger of the rubber hand with a delay of approximately 0.5 seconds by pressing the lever under the rubber hand up (asynchrony). Further, it allowed us to manipulate whether the movement was active or passive by either having the participants lift their index finger up actively, or having the experimenter push the index finger of the participant up by pressing the lever. Finally, this setup allowed us to manipulate the anatomical orientation of the rubber hand by either having the rubber hand placed in an anatomically congruent position, giving the impression of it being continuous with the body, or having the rubber hand being rotated 180 degrees to an anatomically incongruent position (H. H. Ehrsson et al., 2004).

Throughout the experiment, the participants were asked to maintain fixation on the rubber hand. The participants received verbal instructions through the headphones, which consisted of two prerecorded 1-second-long audio clips of either “tap finger” or “relax”. During the active conditions, the participants were asked to do a tapping motion with their right index finger. The tapping motion was done by extending and then flexing the metacarpophalangeal joint while keeping the proximal and distal interphalangeal joints static, in other words, tapping with a straight finger (as in Kilteni & Ehrsson, 2012). In the active conditions the tapping was self-paced, and the participants had to produce a regular rhythm of taps at approximately 1 Hz without the support of a metronome or other external cues. Before the scan started the participants were trained to produce the required tapping movements in a practice trial that lasted a few minutes. In this practice trial the participants listened to a 1 Hz metronome while tapping their right index finger in the moving rubber hand illusion condition, and where then asked to continue tapping without the metronome. Self-paced tapping was chosen to ensure internally generated movement rather “externally triggered” movement (Passingham, 1993), thus avoiding potential interactions between external cue processing and agency. They were also trained to generate the tapping movement with a certain amplitude (3 cm; see further below). If the participants failed to maintain a reasonably consistent tapping frequency or amplitude, they received feedback from the experimenter and did one more practice trial until the participants were consistent and reliable in their tapping frequency and amplitude throughout the trial. In the passive conditions the participants were relaxing their index finger and the experimenter generated the movements as described above. In these passive conditions, the experimenter matched the frequency of the participant’s self-paced movements in the preceding active condition. To make sure that the amplitude of each tap that the experimenter produced was consistent, the experimenter was guided by the measuring stick taped to the table indicating the 3 cm movement amplitude target (see further below). In all conditions, the experimenter was hidden from sight of the participant by standing on the left side of the scanner bore behind the cloth that also occluded the view of the participants real hand (Fig. 1A). The experimenter received continuous instructions about the onset and end of the conditions through headphones as well as through text on a screen that displayed the next condition (the screen was placed in the control room and facing scanner through the glass window of the control room, so that it could be seen from the location of the experimenter inside the scanner room).

### Movement registration and optical sensor

Underneath the index finger of the participant, approximately 3 millimeters proximal to the hole that the rod connected to the plastic ring passed through, there was another small hole (2 mm in diameter). In this smaller hole a fiber optic cable attached to an optical sensor (Omron E3X-HD11, Omron Industrial Automation, Osaka, Japan) was placed, which was able to register when the participant’s index finger was lifted off the table and when it returned to the table during the tapping movements. The optical sensor registered the luminance from the fiberoptic cable with pre-set thresholds so that it recorded dichotomic on/off data (finger on or lifted off) with a sampling frequency of 100 Hz and saved this to a text file. This allowed us to record the frequency of taps, the duration of each tap and the total number of taps in each participant and in each condition. As described above, the experimenter had a measuring stick taped to the table and could visually inspect that the participant’s taps reached the same amplitude of approximately 3 centimeters, ensuring that the amplitude of the taps was consistent across conditions.

### Design

To test the hypothesis that the sense of agency and the sense of body ownership have different neural substrates, and identify possible neural interactions when the two co-occur, we opted for a full factorial design with 2×2×2 conditions (movement type: active/passive; timing: synchronous/asynchronous; and orientation: congruent/incongruent) (Fig. 2, panel A), giving rise to eight unique conditions (Fig. 2, panel B). This factorial design allowed us to isolate neural correlates of the sense of agency and the sense of body ownership (as the two-way interactions) while controlling for basic effects related to differences between active and passive movements, visuo-somatosensory and visuo-motor synchrony, and visual impressions from observing the rubber hand in different orientations (as the three main effects); and also examine possible (three-way) interaction between body ownership and agency when combined in the moving RHI condition. Since the two two-way interactions defining ownership and agency are orthogonal in this design, we can also examine their overlap in activation by using a conjunction analysis. Thus, we reasoned that this experimental design would be ideal to address the questions were interested in.

**Figure 2.**
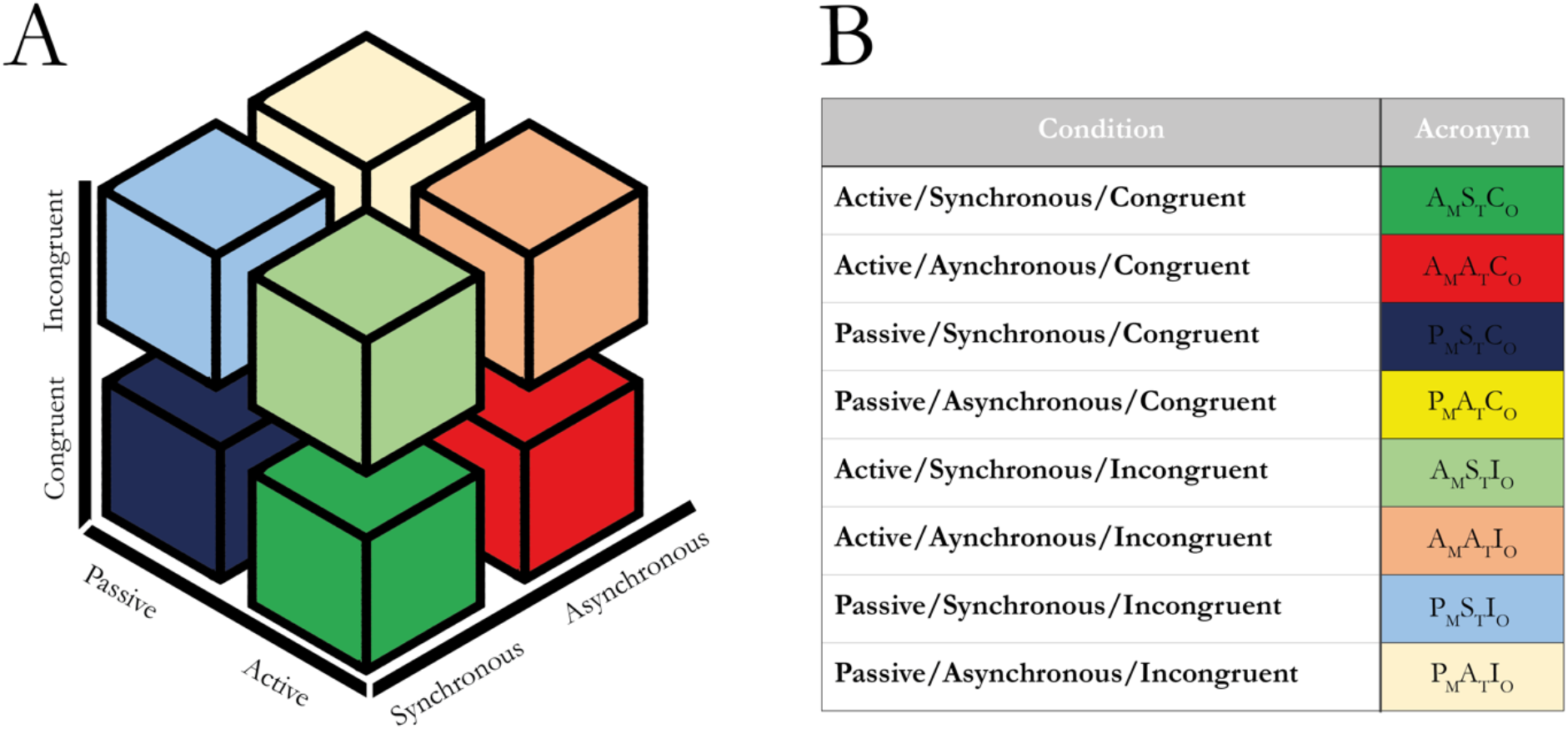
**A** Schematic illustration of the design matrix for the 2×2×2 factorial giving rise to eight unique conditions. **B** All eight unique conditions and their acronyms used in this paper. Each letter indicating the movement type (active or passive), the timing of the movements (synchronous or asynchronous) and the orientation of the rubber hand relative to the participant’s hand (congruent or incongruent) are followed by a subscripted letter indicating which factor the letter belongs to (M=movement type, T=timing, O=orientation).

The fMRI experiment was designed as a block design given the efficiency of this design type (Friston et al., 1999). The experiment was divided into four runs, where two runs were collected with the rubber hand in the congruent position and two runs with the rubber hand in the incongruent position. The separation of the congruent and incongruent trials in separate runs was done due to the fact that it took about a minute to properly reorient the rubber hand, which made it unfeasible to do within a run. The order of the runs was randomized. Each block (epoch) contained a stimulation period of 45 seconds followed by a 5 second resting baseline before the next condition. Each run contained four repetitions of each of the four conditions in said run, totaling in 8 blocks per condition per participant. Every four blocks, there was a 30 second block of a rest baseline condition in which the participants looked at the rubber hand without performing or observing any movement (Fig. 3).

**Figure 3.**
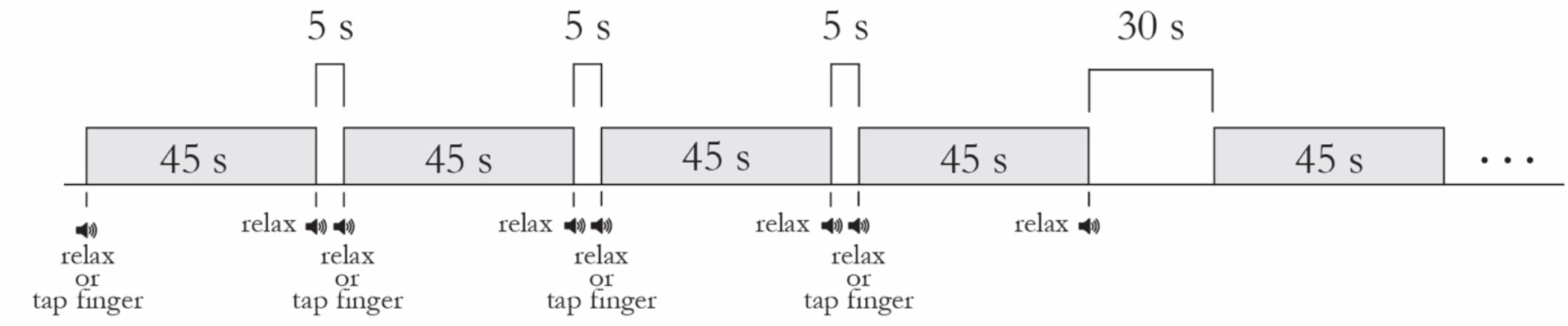
Schematic illustration of the fMRI block design. Each stimuli block consisted of one of the eight conditions with 45 seconds of continuous finger tapping, either actively or passively. Between each block there was a five second rest baseline. After every four blocks there was a longer 30 second rest condition. Four of the eight conditions were repeated four times in each run, since the congruent and incongruent conditions were split in separate runs. The participants received auditory instructions at the beginning and end of each block that consisted of a 1-second-long prerecorded voice saying, “tap finger” or “relax”.

### Behavioral experiment

Prior to the fMRI experiment, all subjects participated in a behavioral experiment. The rationale was fourfold : 1) we wanted to verify that the behavioral paradigm worked as expected for the purpose of the fMRI design; 2) we wanted to quantify ownership and agency using the extensive questionnaires that has been used in previous studies and which are unpractical to use during the scan sessions; 2) since the current eight conditions has never been tested in a single within-subjects design before (Kalckert and Ehrsson 2012 tested the various conditions we use in separate experiments), we also wanted to test for a possible interaction between ownership and agency; 4) we wanted to register how fast the moving RHI was induced in this group of participant exposed to the current paradigm, in order to take this into account in the later fMRI analysis.

Thus, in this behavioral experiment, the participants were tested with the identical moving rubber hand illusion setup that would be used in the MR scanner but laying on a bed in our behavioral testing lab instead. The position of the participants limbs, head and body was the same as during the MR scans. The participants had all eight conditions repeated once and received a 16-statement questionnaire at the end of each condition that probed the illusory experience of the sense of body ownership and the sense of agency (table 1; based on Kilteni & Ehrsson, 2012). Control questions probing suggestibility and task compliance were also included. The questionnaire was rated on a 7-point Likert scale ranging from (−3) to (+3), with (−3) corresponding to “completely disagree”, (+3) corresponding to “completely agree”, and (0) corresponding to “neither agree nor disagree”. The stimulation period for each condition was 45 seconds. When all conditions had been tested, six more trials, three with the active/synchronous/congruent (A_M_S_T_C_O_) condition and three with the passive/synchronous/congruent (P_M_S_T_C_O_) condition were conducted. In these additional trials of the A_M_S_T_C_O_ and P_M_S_T_C_O_ condition, the illusion was induced in the same manner, but this time the participants were instructed to verbally indicate when they started to experience that the “rubber hand was their hand” (corresponding to the fourth statement in the ownership questionnaire, table 1; (H. H. Ehrsson et al., 2004)). This yielded average “illusion onset time” measurements for each participant in both the A_M_S_T_C_O_ and P_M_S_T_C_O_ conditions (Supplementary Table 1). These time individual intervals (A_M_S_T_C_O_ range 0-30 s, mean 11.5 ± 8.2 s, P_M_S_T_C_O_ range 0-30.2, mean 12.26 ± 9.1 s, non-significant difference between onset time in A_M_S_T_C_O_ and P_M_S_T_C_O_, W=127.00, p=0.346, Rank-Biserial Correlation -0.218) were then used to define the start of the illusion conditions of interest in the fMRI analyses (see further below). This allowed us to focus on our analysis on the periods when the moving rubber hand illusion had been elicited (H. H. Ehrsson et al., 2004). The periods before the illusion onset times were modelled as conditions of no interest and not used in the statistical contrasts.

**Table 1.** The statements used in the questionnaire experiment conducted before the fMRI study (the “behavioral pre-test”). Each statement was rated once per condition. The statements were rated on a 7-point Likert scale ranging from (−3) to (3). There were four statements assessing the sense of body ownership and the sense of agency respectively, as well as four control statements for both the sense of body ownership and the sense of agency, respectively.

| Assessment | Statement |
| --- | --- |
| Ownership | I felt as if I was looking at my own hand |
|  | I felt as if the rubber hand was part of my body |
|  | It seemed as if I were sensing the movement of my finger in the location where the rubber finger moved |
|  | I felt as if the rubber hand was my hand |
| Agency | The rubber hand moved just like I wanted it to, as if it was obeying my will |
|  | I felt as if I was controlling the movements of the rubber hand |
|  | I felt as if I was causing the movement I saw |
|  | Whenever I moved my finger I expected the rubber finger to move in the same way |
| Ownership control | I felt as if my real hand were turning rubbery |
|  | It seems as if I had more than one right hand |
|  | It appeared as if the rubber hand were drifting towards my real hand |
|  | It felt as if I had no longer a right hand, as if my right hand had disappeared |
| Agency control | I felt as if the rubber hand was controlling my will |
|  | I felt as if the rubber hand was controlling my movements |
|  | I could sense the movement from somewhere between my real hand and the rubber hand |
|  | It seemed as if the rubber hand had a will of its own |

### fMRI data acquisition

The experiment was conducted using a 3 Tesla GE MR750 scanner equipped with an 8-channel head coil. T2*-weighted gradient echo EPIs with BOLD contrast were used as an index of brain activity (Logothetis, 2003; Logothetis et al., 2001). Each functional volume consisted of 43 continuous slices with 3 mm slice thickness and 0.5 mm interslice space. The field of view (FOV) was defined as a matrix with the dimensions 72×72 (3×3 mm in plane resolution, TE=30 ms) ensuring coverage of the whole brain. One volume was collected every 2.048 s (TR=2048 ms) and a total of 1812 functional volumes were collected for each participant, divided into 4 runs of 453 volumes each. A high-resolution structural image was collected for each participant at the end of the experiment (3D MPRAGE sequence, 1×1×1 mm voxel size, FOV 240 × 240 mm, 180 slices, TR=6404 ms, TE=2808 ms, flip angle = 12°).

### Statistical analysis

#### Questionnaire data

The data from the behavioral pre-test experiment was tested for normality using the Shapiro-Wilk’s test. If the data deviated from normality, the results were analyzed using non-parametric Wilcoxon Signed Ranks Test. The questionnaire data from the pre-testing was analyzed using JASP (version 0.11.1, 2019, University of Amsterdam, The Netherlands). For each participant, the subjective ratings from the four statements probing ownership was averaged into a ownership score, the four agency statements into an agency score, and the control statements were similarly averaged into an ownership control score and agency control score, respectively (Kilteni & Ehrsson, 2012). For each condition, a sense of body ownership or agency was defined as a mean ownership or agency score of >0. To test for body ownership or agency within a condition and control for unspecific suggestibility effects, the ownership score was compared statistically to the ownership control score and the agency score to the agency control score, respectively. To compare body ownership and agency between conditions, an ownership index and agency index was calculated. The indices were defined as the difference between the ownership score and the ownership control score (ownership index), and between the agency score and agency control score (agency index), respectively (Abdulkarim & Ehrsson, 2016).

#### Movement sensor data

The data from the optical sensor was analyzed using MatLab (version 2018b, statistical toolbox, Mathworks, Massachusetts, USA). The optical sensor was not available for the first 10 participants (still under development due to unexpected delay), which is why we only included data from the optical sensor from 19 participants. The number of taps from each trial was extracted for participant 10-29. The frequency of taps was calculated by dividing the number of taps by each condition’s total time. The number of taps as well as the frequency of taps was then averaged across participants for each condition. The statistical analysis focused on testing for main effects of synchrony, active or passive movements and congruent or incongruent rubber hand orientation in terms of the number and frequency of taps in line with the fMRI design.

#### fMRI data preprocessing, modeling and statistical inference

The fMRI data from all participants was analyzed using Statistical Parametric Mapping 12 (SPM12; Wellcome Trust Center for Neuroimaging, University College London, UK). Before the functional imaging data underwent the preprocessing steps, all functional and anatomical images were rotated back to the standard position, which they deviated from due to the forward head tilt inside the scanner coil. After that, the preprocessing steps including motion correction, slice timing correction, co-registration, normalization (to the Montreal Neurological Institute (MNI) standard brain). The functional images were resampled to a resolution of 2×2×2 mm, and the spatial smoothing was applied using a 6 mm FWHM Gaussian kernel. The statistical analysis was done by fitting a general linear model (GLM) to the data for each participant. The hemodynamic response function was convolved with boxcar regressors for each condition of interest. Linear contrasts were defined at the individual level and exported to the second level random-effects analysis. Importantly, we modeled the first period in each condition as a condition of no interest, based on the time it took for each individual participant to experience the illusion in the behavioral pre-test (see above), and the periods from illusion onset to the end of each condition as the condition of interest used in our main analyses (in line with (H. H. Ehrsson et al., 2004; Guterstam et al., 2013). For the A_M_S_T_C_O_ and P_M_S_T_C_O_ conditions, we used their corresponding rubber hand illusion onset times, whereas for all other conditions (that did not trigger the rubber hand illusion), we used the average of the A_M_S_T_C_O_ and P_M_S_T_C_O_ times.

For the main contrasts, we had anatomical hypotheses regarding which regions we expected to be activated during experiences of body ownership and agency based on the previous fMRI literature (see introduction), and therefore in these regions report the results that are statistically significant at a threshold of p<0.05 after small volume correction (family-wise error correction; “FWE”). However, since earlier ownership studies used brushstrokes or similar tactile stimulation applied to relaxed hands instead of finger movements, we anticipated that the exact location of peaks associated with the rubber hand illusion could change within the hypothesized frontal, parietal, and subcortical regions. Therefore, the volumes of interest used in the small volume correction were centered on peak coordinates obtained from a “localizer” study where we used the same 3T MR-scanner and fMRI scanning protocol as in the main experiment to identify the locations of active candidate areas during the moving rubber hand illusion. In brief, the localizer study included 27 participants looking at and controlling the index finger movement of a robotic hand wearing a plastic glove identical in shape and size to the rubber hand used in our current experiment. This robotic “rubber hand” was placed in view of the participant on a supporting table in a very similar arrangement to the one used in the current study. When the participant moved his or her index finger the rubber hand moved its index finger in the same way and synchronously, triggering the moving rubber hand illusion (verified with illusion questionnaire ratings which were affirmative in most participants; data not shown). In the localizer study we contrasted this illusion condition (corresponding to the A_M_S_T_C_O_ condition in the present study) to a resting baseline condition where the participants were just looking at the rubber hand without performing or observing any movement. Peaks from this localizer contrast were then used to define the coordinates in MNI space for the spheres (10 mm in radius) in the small volume corrections (Supplementary Table 2 for list of all peaks used from this localizer study to define the volumes of interest). For the intraparietal cortex – a region often associated with the RHI and illusory hand ownership in the previous fMRI literature – we added peaks from (H. H. Ehrsson et al., 2004) since no activations were detected in the localizer contrast in this region. Similarly, for left insular cortex and right angular gyrus in the temporo-parietal junction (TPJ) region – two areas often associated with different aspects of agency in the previous literature – we used coordinates from classic neuroimaging agency studies (Farrer et al., 2003; Farrer & Frith, 2002) to define peaks for small volume correction in these regions. In the rest of the brain, i.e. outside the regions related to our a priori define anatomical hypotheses, we corrected for the number of comparisons in the whole brain space using a test of false discovery rate (FDR) set at p<0.05. All our statistical inferences and main findings are based on results that survive multiple comparison correction based on these two approaches, which collectively balances type 1 and type 2 errors and hypothesis-driven and explorative approaches.

**Table 2.** Activation peaks for the main contrasts. **A**. The sense of body ownership in the moving rubber hand illusion expressed as the interaction between synchrony and orientational congruency between the participant’s real hand and the rubber hand, defined as the contrast [(P_M_S_T_C_O_–P_M_A_T_C_O_)–(P_M_S_T_I_O_–P_M_A_T_I_O_)] + [(A_M_S_T_C_O_–A_M_A_T_C_O_)– (A_M_S_T_I_O_–A_M_A_T_I_O_)] **B**. The sense of agency expressed as the interaction between synchrony and movement type (active/passive), defined as the contrast [(A_M_S_T_C_O_–P_M_S_T_C_O_)–(A_M_A_T_C_O_–P_M_A_T_C_O_)] + [(A_M_S_T_I_O_–P_M_S_T_I_O_)–(A_M_A_T_I_O_– P_M_A_T_I_O_)]. **C**. The three-way interaction between synchrony, movement type (active/passive) and orientation, representing the areas that demonstrate increased activity when experiencing agency over bodily objects as opposed to external objects, defined as the contrast [(A_M_S_T_C_O_–P_M_S_T_C_O_)–(A_M_A_T_C_O_–P_M_A_T_C_O_)]–[(A_M_S_T_I_O_–P_M_S_T_I_O_)– (A_M_A_T_I_O_–P_M_A_T_I_O_)]. **D**. The inverse of the three-way interaction between synchrony, movement type (active/passive) and orientation, representing the areas that demonstrate increased activity when experiencing agency over external objects as opposed to bodily objects, defined as the contrast [(A_M_S_T_C_O_–P_M_S_T_C_O_)–(A_M_A_T_C_O_– P_M_A_T_C_O_)]–[(A_M_S_T_I_O_–P_M_S_T_I_O_)–(A_M_A_T_I_O_–P_M_A_T_I_O_)]. PrCG= precentral gyrus, PoCG= postcentral gyrus, SMG= suprmarginal gyrus, ITG= inferior temporal gyrus, dlFPC= dorsolateral prefrontal cortex, mPFC= medial prefrontal cortex, IPS= intraparietal sulcus, IOG= inferior occipital gryus, STG= superior temporal gyrus, MOG= middle occipital gyrus. *\* indicates activation peaks that survive small volume correction (FWE correction, p<0*.*05 corrected);* ^*+*^ *indicates an activation peak in the agency contrast that almost reached statistical significance after small volume correction (FWE correction); the peaks without an asterix did not survive small volume correction and are reported with their uncorrected p-value*.

| Anatomical region | MNI *x,y,z* (mm) | Peak *t* | *p*-value |
| --- | --- | --- | --- |
| L PrCG (PMd) | -42, -10, 58 | 4.66 | 0.009** |
| L PrCG (PMd) | -34, -10, 64 | 4.30 | 0.019* |
| L PrCG (PMd) | -42, -12, 56 | 4.13 | 0.010* |
| L PrCG (PMd) | -36, -10, 62 | 3.99 | 0.014* |
| L SMG | -60, -48, 38 | 3.69 | 0.025* |
| R Cerebellum (lobule VIIb) | 4, -68, -46 | 3.47 | 0.038* |
| R Cerebellum (lobule VIIa) | 40, -74, -34 | 3.28 | 0.027* |
| R Cerebellum (lobule VIIa) | 38, -72, -24 | 3.19 | 0.034* |
| R ITG | 42, -70, -8 | 3.38 | 0.046* |
| L dlPFC | -24, 42, 38 | 4.42 | 0.001 |
| R mPFC | 6, 50, 40 | 3.33 | 0.001 |
| L PrCG (M1) | -30, -22, 62 | 3.21 | 0.001 |
| L PoCG (S1) | -36, -22, 50 | 3.72 | <0.001 |
| L PoCG | -46, -14, 52 | 3.51 | <0.001 |
| L PoCG | -36, -32, 66 | 5.01 | <0.001 |
| L PoCG | -56, -16, 40 | 3.05 | 0.003 |
| L IPS | -26, -76, 42 | 4.91 | <0.001 |
| R IOG | 44, -70, -12 | 2.97 | 0.003 |

**B**
| Anatomical region | MNI *x,y,z* (mm) | Peak *t* | *p*-value |
| --- | --- | --- | --- |
| R STG | 58, -24, 12 | 4.03 | 0.051<sup>+</sup> |
| L PrCG (PMd) | -38, -8, 62 | 3.89 | 0.013* |
| R STG | 60, -20, 12 | 5.12 | <0.001 |
| L STG | -50, -28, 6 | 4.88 | <0.001 |
| L PoCG (S1) | -52, -30, 54 | 3.51 | 0.001 |
| R IPS | 36, -40, 52 | 3.47 | 0.001 |
| L IPS | -36, -40, 46 | 3.69 | <0.001 |

**C**
| Anatomical region | MNI *x,y,z* (mm) | Peak *t* | *p*-value |
| --- | --- | --- | --- |
| L PoCG (S1) | -38, -28, 52 | 4.21 | 0.007* |
| L PoCG (S1) | -36, -26, 38 | 2.85 | 0.004 |
| L PoCG | -54, -18, 28 | 2.96 | 0.003 |
| L PoCG | -52, -16, 12 | 3.26 | 0.001 |

**D**
| Anatomical region | MNI *x,y,z* (mm) | Peak *t* | *p*-value |
| --- | --- | --- | --- |
| L MOG | -20, -94, 0 | 4.08 | <0.001 |
| R MOG | 26, -92, 4 | 3.08 | 0.002 |

Some activations that did not survive correction for multiple comparisons are still mentioned in the text or shown in figures as part of the statistical parametric maps produced by SPM12 (based on a threshold of p<0.005 uncorrected). We report these for purely descriptive purposes (Gentile et al., 2013; Preston & Ehrsson, 2016) and always clearly identify these as not reaching our significance criterion. We report these non-significant activations mainly for five reasons: (i) false negatives and limited sensitivity is a concern in fMRI studies, so being overly conservative might conceal potentially interesting results (ii) we want to report the activation maps in a transparent fashion and not only describe those regions that were part of our hypothesis; (iii) activation peaks that did not survive correction for multiple comparison can still be used to define anatomical hypotheses for future fMRI studies; and (iv) the reporting of the entire activation maps including non-significant activation can provide information about the anatomical specificity of these latter effects (i.e. single active brain area or widespread effects in many regions); (v) non-significant peaks can be used in future imaging meta-analysis where it is often important to have data from the whole brain (and not only a few peaks that survive multiple comparisons correction). As mentioned, all main conclusions in the manuscript are based on activations that are significant (in one case almost significant) after correction for multiple comparisons, i.e. p<0.05 after FWE correction.

The visualization of the results is done by overlaying the peaks on an 3D rendering of a standard MNI brain using Surf Ice (https://www.nitrc.org/projects/surfice/), as well as on sections from the average anatomical image for all participants. The anatomical localizations of the activations based on macroanatomical landmarks (sulci and gyri) using the terminology from the Duvernoy and Parratte brain atlas (Duvernoy, 1999). In addition, we have used the SPM Anatomy toolbox v2.2 to relate somatosensory activations to population-based cytoarchitectonic data (Eickhoff et al., 2005). All coordinates for the activation peaks are given in MNI space. Contrast estimates for each significant peak were extracted using MatLab (version 2018b) and presented in bar charts together with the corresponding standard errors (SEs) for purely descriptive purposes. In line with the SPM approach, we make no further statistical analyses on these bar chart plots, but all conclusions and statistical inferences are based on significant (two-way and three-way) interaction contrasts in line with our factorial design.

### Planned fMRI analyses

To identify regions that display BOLD responses that reflect the sense of body ownership or the sense of agency we defined linear contrasts that corresponded to the two-way interactions that captured ownership and agency in our factorial design (ownership: interaction timing x orientation; agency: interaction timing x movement type). In other words, we subtracted the control conditions where no illusory experience in question was present (or strongly suppressed) from the experimental condition in which they were present. Thus, for the sense of body ownership, we defined the contrast [(P_M_S_T_C_O_– P_M_A_T_C_O_)–(P_M_S_T_I_O_–P_M_A_T_I_O_)] + [(A_M_S_T_C_O_–A_M_A_T_C_O_)–(A_M_S_T_I_O_–A_M_A_T_I_O_)], including both the active and passive conditions. This contrast corresponds to the interaction between the factors synchrony (of the movements) and congruency (between the orientation of the rubber hand with the participant’s real hand) since we know asynchronous movements and anatomical incongruency to abolish the sense of ownership of the rubber hand in the moving rubber hand illusion (Kilteni & Ehrsson, 2012). Similarly, for the sense of agency, we defined the contrast [(A_M_S_T_C_O_–P_M_S_T_C_O_)–(A_M_A_T_C_O_–P_M_A_T_C_O_)] + [(A_M_S_T_I_O_– P_M_S_T_I_O_)–(A_M_A_T_I_O_–P_M_A_T_I_O_)], including both the congruent and incongruent conditions. This contrast is the interaction between the two factors timing (synchronous or asynchronous) and type of movement (active or passive), because we know that both the sense of volition associated with active movements and the match between expected and actual sensory feedback from the movements are required in order for a sense of agency to develop (Haggard, 2017); hence both asynchronous movements and passive movements should abolish the sense of agency of the rubber hand. Note, that these key contrasts are balanced and fully matched in terms of magnitude of visual and somatosensory stimulation related to the observed and felt movements, as well as frequency and amplitude of finger taps (Supplementary table 3), and thus isolates the neural activities related to ownership and agency we are interested in.

A further strength of this design is that the two interaction contrasts that operationalize ownership and agency are orthogonal (i.e., independent), which means that we can also test for active voxels that are significantly active in both contrasts by using a conjunction analysis. Thus, this conjunction analysis identifies active areas that show increases in activity that reflect both ownership and agency.

Finally, and not the least, the current 2×2×2 factorial design allow us to investigate the interaction between the sense of body ownership and the sense of agency. To this end, we defined a linear contrast that was composed as a three-way interaction between the three factors in the factorial design, namely movement type, synchrony and orientation of the fake hand, congruent or incongruent with the real hand. This contrast [(A_M_S_T_C_O_–P_M_S_T_C_O_)–(A_M_A_T_C_O_–P_M_A_T_C_O_)]–[(A_M_S_T_I_O_–P_M_S_T_I_O_)–(A_M_A_T_I_O_–P_M_A_T_I_O_)] identifies a neural response that specifically reflect the combination of body ownership and agency when voluntary moving one’s body. This can reflect stronger sense of ownership during active movements or differences in agency over an own body part (the rubber hand during the rubber hand illusion) as compared to agency an external object (the rubber hand in the incongruent orientation that does not feel like part of one’s body).

### Post-hoc fMRI connectivity analyses

Task-related connectivity was assessed by performing a psycho-physiological interaction (PPI) analysis. The PPI indexes task or contrast specific changes in the connectivity between two brain regions. A significant PPI that indicates that the correlation of the brain activity in the two regions (measured as the change in the slope of their linear regression curve) changes significantly with the experimental or psychological context (Friston et al., 1997). To follow up the regional results (see below) we decided to conduct a post-hoc PPI analysis for purely descriptive purposes. We placed a seed voxel in the post-central gyrus contralateral to the stimulated hand. The seed was selected based on activity in this region that was elucidated during the three-way interaction contrast described above. The seed was defined for each participant as a 10 mm sphere around the group level activation in the post-central gyrus. From this, the time series of activity (first eigenvariate) was extracted and entered into the PPI analysis with the contrast weights from the three-way interaction. The PPI regressors created at the individual level were analyzed at the group level using one-sample t-tests.

### Post-hoc descriptive correlation analysis of ownership contrast and questionnaire ratings

In a post-hoc complementary approach we explore a possible relationship between the subjective ratings of ownership as rated by the individual participants in the questionnaires in the behavioral experiment (before the fMRI) and the contrast that describe the ownership-related activation. Unlike agency that can be experienced by everybody, the feeling of ownership in the rubber hand illusion is vividly experienced in approximately 60-80 % of participants (H. H. Ehrsson et al., 2005; Kalckert & Ehrsson, 2014; Lloyd, 2007) making it possible to probe how individual differences in illusion strength relate to brain activation. Previous studies have shown such a relationship in the premotor cortex (H. H. Ehrsson et al., 2004, 2005; Gentile et al., 2013). To this end we conducted analyses of the fMRI data combined with a behavioral covariate. For each participant, we calculated “behavioral contrast” (analogous to the defined contrasts in the fMRI analyses; [(P_M_S_T_C_O_–P_M_A_T_C_O_)–(P_M_S_T_I_O_–P_M_A_T_I_O_)] + [(A_M_S_T_C_O_–A_M_A_T_C_O_)–(A_M_S_T_I_O_–A_M_A_T_I_O_)]) of the ownership ratings from all eight conditions and entered this as a covariate in the GLM model in the second level analysis together with the contrast images reflecting the ownership contrast. This analysis allowed us to examine whether stronger subjective ownership in the synchronous and congruent conditions (A_M_S_T_C_O_ and P_M_S_T_C_O_ compared to the other conditions) correlated with stronger BOLD signals specifically in the ownership contrast.

### Post-hoc conjunction analysis across ownership and agency contrasts

To investigate which brain regions that showed an overlapping activation in both the ownership and agency contrasts, we conducted a conjunction by performing a one-way ANOVA at the second level analysis and entering the two different first level contrasts as the groups in the one-way ANOVA. The contrasts are then specified for the two groups and combined as an inclusively mask.

## Results

### Behavioral experiment

The results from the behavioral pre-test experiment replicated the main findings from the original moving rubber hand illusion paper and confirmed that our behavioral paradigm worked as expected (Kilteni & Ehrsson, 2012), but in a full 2×2×2 factorial within-subject design. The results confirmed that the sense of body ownership and sense of agency can be dissociated behaviorally as we had expected (Fig. 4, Panel A). In the A_M_S_T_C_O_ conditions, the participants experienced both a sense of body ownership and agency of the rubber hand, i.e., the mean rating scores of these two sensations were both positive meaning that the participants on average affirmed both these experiences in the classic moving RHI condition with active finger movements. Further, in the P_M_S_T_C_O_ condition, the classic moving RHI condition with passive finger movements, the participants experienced a sense of body ownership (positive rating score) of the rubber hand but denied sensing of agency (negative mean agency score). Finally, in the A_M_S_T_I_O_ condition the participants experience a sense of agency of the rubber hand but no sense of body ownership (positive and negative agency rating scores, respectively). In the control conditions the participants did not report sensing body ownership or agency and the mean ownership and agency scores were negative (Fig. 4, Panel A).

**Figure 4.**
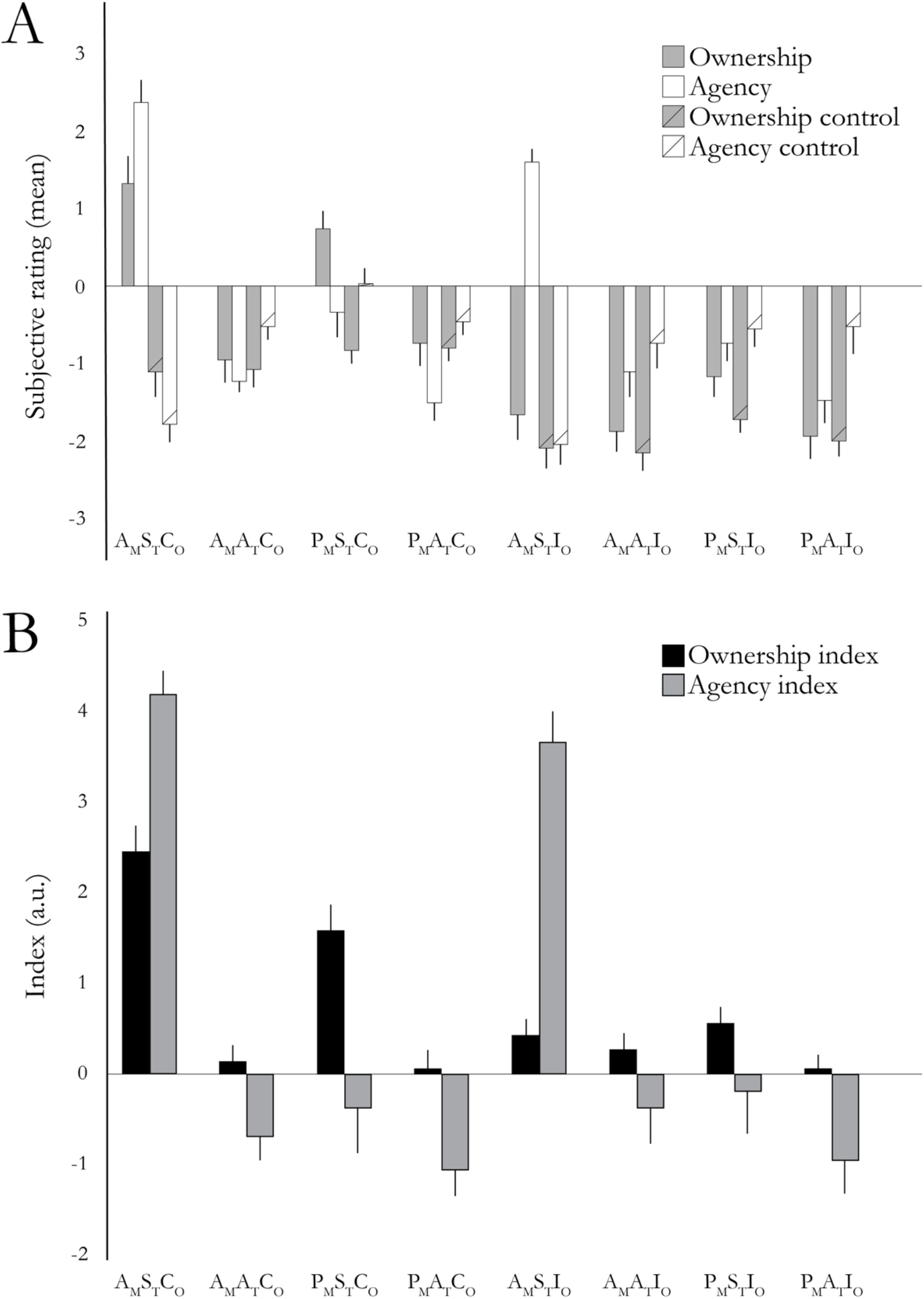
**A**. The results from the behavioral pre-test. These results show a double dissociation between the sense of body ownership and sense of agency in our full factorial design. The A_M_S_T_C_O_ condition displayed high ratings for both sense of body ownership and sense of agency. The P_M_S_T_C_O_ condition showed high ownership ratings and low agency ratings, whereas the A_M_S_T_I_O_ condition showed high agency ratings and low ownership ratings. Bars represent mean ratings and error bars indicate the SEM. **B**. Ownership and agency indices calculated by subtracting the pooled ownership and agency control ratings from the pooled ownership and agency ratings, respectively. Bars indicate the means and error bars the SEM.

We then compared the ownership to the ownership control ratings and found significantly higher ratings of the ownership statements compared to the control statements in the A_M_S_T_C_O_ condition (W=349, p<0.001, Rank-Biserial correlation 0.989) and P_M_S_T_C_O_ condition (W=314, p<0.001, Rank-Biserial correlation 0.932). The same analysis for the sense of agency showed significantly higher ratings of the agency statements compared to the agency control statements in the A_M_S_T_C_O_ condition (W=351, p<0.001, Rank-Biserial correlation 1.00) and A_M_S_T_I_O_ condition (W=351, p<0.001, Rank-Biserial correlation 1.00). The individual ratings for each statement and condition are given in (Supplementary table 4).

We then directly tested the hypothesis that the sense of body ownership depended on synchronous visuo-somatosensory feedback when moving the finger as well as anatomical congruency between the rubber hand and the participants real hand (Botvinick & Cohen, 1998; H. Ehrsson, 2012; Guterstam, Larsson, et al., 2019; Tsakiris, 2010). To this end we analyzed the ownership indices (the difference between the ownership score and ownership control score to control for suggestibly effects) in a 2×2×2 ANOVA (see Fig. 4, Panel B for ownership and agency indices across the eight conditions). The factors *movement* type (active/passive), *timing* of movements (synchronous/asynchronous), and *orientation* of the rubber hand (congruent/incongruent) were entered in the analysis. The results showed a significant main effect of movement (F=6.63, df 29, 1, p=0.016), significant main effect of timing (F=41.276, df 29, 1, p<0.001), and significant main effect of orientation (F=17.645, df 29, 1, p<0.001). Importantly, the interaction between timing and orientation was significant (F=31.933, df 29, 1, p<0.001), line with our operationalization of ownership in the fMRI factorial experimental design. There was no significant interaction between timing synchrony and movement type (F=0.894, df 29, 1, p=0.353). However, the interaction between movement type and orientation was also significant (F=5.982, df 29, 1, p=0.022), which suggests higher ownership ratings during the active finger movements when the rubber hand was in an anatomically congruent position. Moreover, there was a significant three-way interaction between timing, movement type and orientation (F=6.421, df 29, 1, p=0.018). This three-way interaction suggests enhanced ownership of the rubber hand in the active moving RHI condition when participants experience both ownership and agency over the moving rubber hand and provide behavioral support for examining the interaction of ownership and agency in our factorial fMRI design. This is in itself a novel behavioral finding that suggest that active finger movements provide a stronger cue for body ownership than passive ones, which goes beyond our previous studies where such an effect could not be assessed (Kilteni & Ehrsson, 2012, 2014, 2017).

Similarly, we hypothesized that the sense of agency is dependent on synchronous visuo-motor feedback when moving the finger (the match between predicted sensory consequences of the active movement) as well as on participants actively moving the index finger (volition associated with generating active movements as such) (Kilteni & Ehrsson, 2012, 2014). To this end we also analyzed the agency indices (the difference between the agency scores and the agency control scores to control for suggestibility effects) in a 2×2×2 ANOVA. The three factors movement type (active/passive), timing (synchronous/asynchronous), and orientation (congruent/incongruent) were entered in the analysis (Fig. 4, Panel B). As expected, the results showed a significant main effect of movement type (F=42.244, df 29, 1, p<0.001) and a significant a main effect of synchrony (F=107.572, df 29, 1, p<0.001), which suggest that both active movements and synchronous seen and felt movement enhanced agency ratings. There was no main effect of orientation (F=0.021, df 29, 1, p=0.886) indicating that the orientation of rubber hand did not influence agency. Importantly, the interaction between synchrony and movement type was significant (F=36.751, df 29, 1, p<0.001) in line with our operationalization of agency as this two-way interaction in our fMRI design. The interaction between movement type and orientation was not significant (F=0.406, df 29, 1, p=0.530) and neither was the interaction between synchrony and orientation (F=0.379, df 29, 1, p=0.251). The three-way interaction between synchrony, movement type and orientation was also non-significant (F=1.560, df 29, 1, p=0.223). Post-hoc pairwise comparisons between the A_M_S_T_C_O_ and P_M_S_T_C_O_ conditions in terms of ownership index, and ownership scores, respectively, revealed significant differences in both cases (t=3.155, df=29, p=0.004; t=2.413, df=29, p=0.023). These latter results are consistent with the hypothesis that agency does not depend on the orientation of the rubber hand but can be flexibly experienced for both body parts as well external events arising as a consequence of bodily movements, and that agency can be operationalized as and interaction between movement type and temporal congruence and only arising for active movements with synchronous visual feedback. Overall, the questionnaire results from our behavioral experiment confirmed that our selective manipulation of ownership and agency in the moving rubber hand illusion worked as expected, and also provided novel behavioral support for an interaction of ownership and agency.

### fMRI

#### The sense of body ownership is associated with activity in multisensory frontal and parietal regions as well as cerebellar regions

To identify activations associated with the sense of ownership of the rubber hand in both the active and passive condition, we used the contrast [(P_M_S_T_C_O_–P_M_A_T_C_O_)–(P_M_S_T_I_O_–P_M_A_T_I_O_)] + [(A_M_S_T_C_O_–A_M_A_T_C_O_)– (A_M_S_T_I_O_–A_M_A_T_I_O_)]. In line with our hypothesis, this contrast revealed significant activation peaks in the left premotor cortex, posterior parietal cortex and cerebellum (p<0.05 FWE corrected for multiple comparisons; Fig. 5; Table 2). The premotor activations were located in the precentral gyrus at a location that corresponds to the dorsal premotor cortex (PMd; -34, -10, 64; p<0.05, FWE corrected; Fig. 5) and in the parietal lobe activations were observed in the supramarginal gyrus (SMG; -60, -48, 38; p<0.05, FWE corrected; Fig. 5). Activation peaks were also observed in the primary motor cortex (precentral gyrus) and the primary somatosensory cortex (postcentral gyrus), at sites that corresponded very well to peaks identified in the localizer experiment (see above). However, since no a priori hypotheses existed for these regions and they did not survive correction for multiple comparisons at the whole-brain level, they are reported with their uncorrected p-values. We also observed activity in the intraparietal cortex (p<0.001 uncorrected), but more posteriorly than we had predicted based on previous work. In the subcortical structures, we observed significant activity in the crus 1 (lobule VIIa; 40, -74, -34) and vermis (lobule VIIb; 4, -68, -46) of the cerebellum (p<0.05, FWE corrected; Fig. 5). Finally, we observed a large active cluster in the left dorsolateral prefrontal cortex (that can be seen in Figure 5, Panel A; p<0.001 uncorrected)). No clusters survived correction for multiple comparisons at the whole-brain level. Further statistical details on the anatomical locations in MNI-space of the above-mentioned peaks are given in Fig. 5 and Table 2.

**Figure 5.**
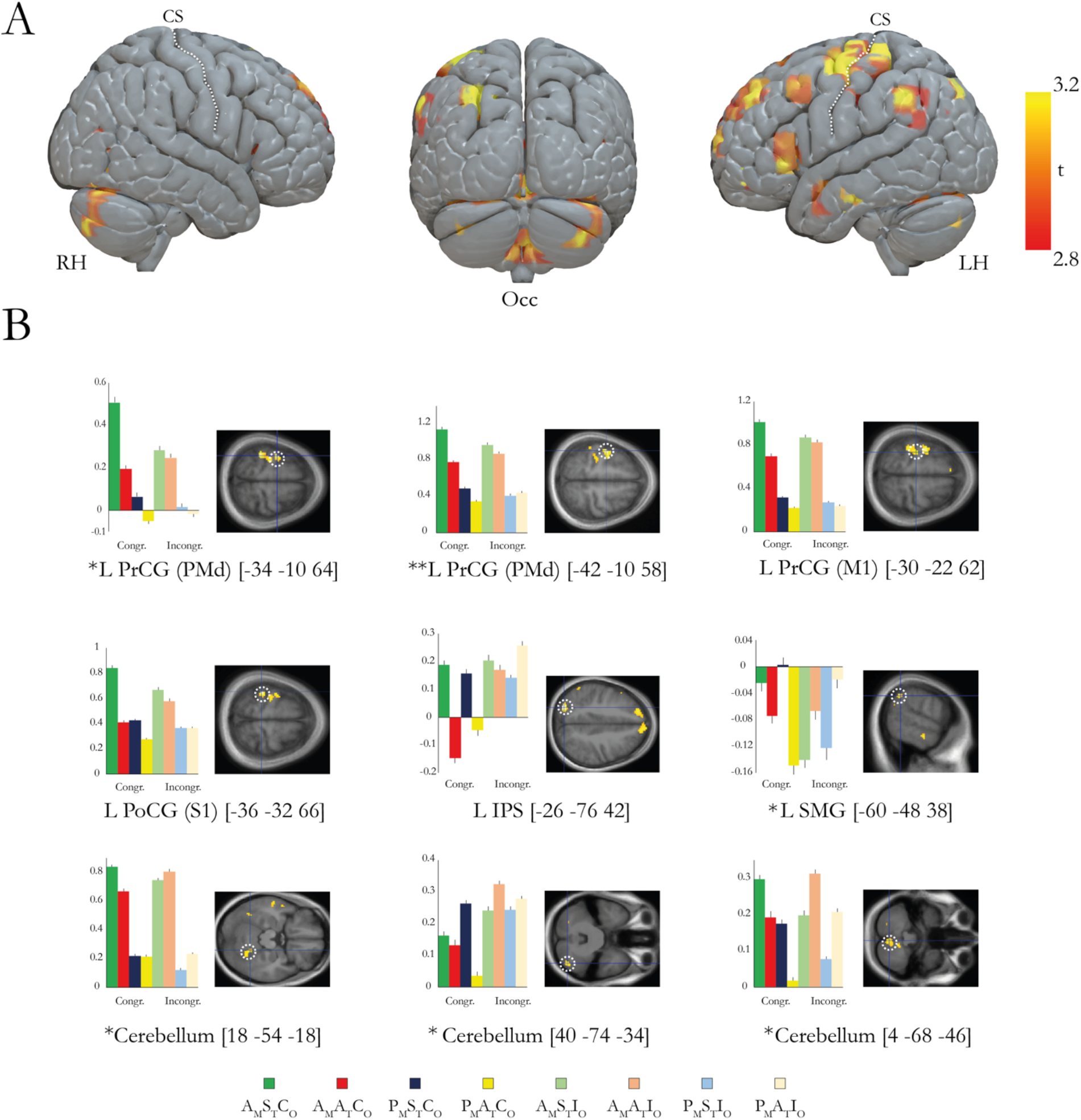
**A**. Overview of the brain regions that display activation reflecting the sense of body ownership over the rubber hand. For display purposes only, the activations are projected on to a three-dimensional render of a standard brain with at a threshold of p<0.005 (uncorrected for multiple comparisons, k≥ 5). RH/LH, right/left hemisphere. Occ, occipital view. CS, central sulcus. **B**. Bar charts displaying the parameter estimates (a.u.) and SEs for the major peaks of activation. The coordinates are given in MNI space. The peaks are displayed in representative sections indicated by a dotted white circle on an activation map (p<0.005 uncorrected for display purposes). L/R, left/right. PrCG, precentral gyrus. PoCG, postcentral gyrus. SMG, supramarginal gyrus. IPS, intraparietal sulcus. Asterixes indicate activation peaks that survive small volume correction (*p<0.05 corrected, ** p<0.01); the peaks without an astrerix did not survive small volume correction and are reported in table 2 with their uncorrected p-value.

#### Correlation between subjective ownership ratings and ownership contrast

In a complementary descriptive approach, we followed up the above ownership-interaction contrast by examining if those BOLD effects also correlated with the subjective ratings in the ownership statements. To this end we performed a multiple regression analysis using the ownership ratings from each participant to search for voxels whose parameter estimates could be predicted from the behavioral contrast (see methods). We identified four such regions whose parameter estimates were significantly correlated with the behavioral contrast (Fig. 6). The activity in in the left premotor cortex (PMd; -24, -12, 70; p<0.05, Fig. 6) and cerebellum was significant after FWE-correction (Cerebellum; -26, -46, -26; p<0.05, Fig. 6) whereas the activity in the postcentral gyrus and postcentral sulcus were not (p<0.001, uncorrected; Fig.6).

**Figure 6.**
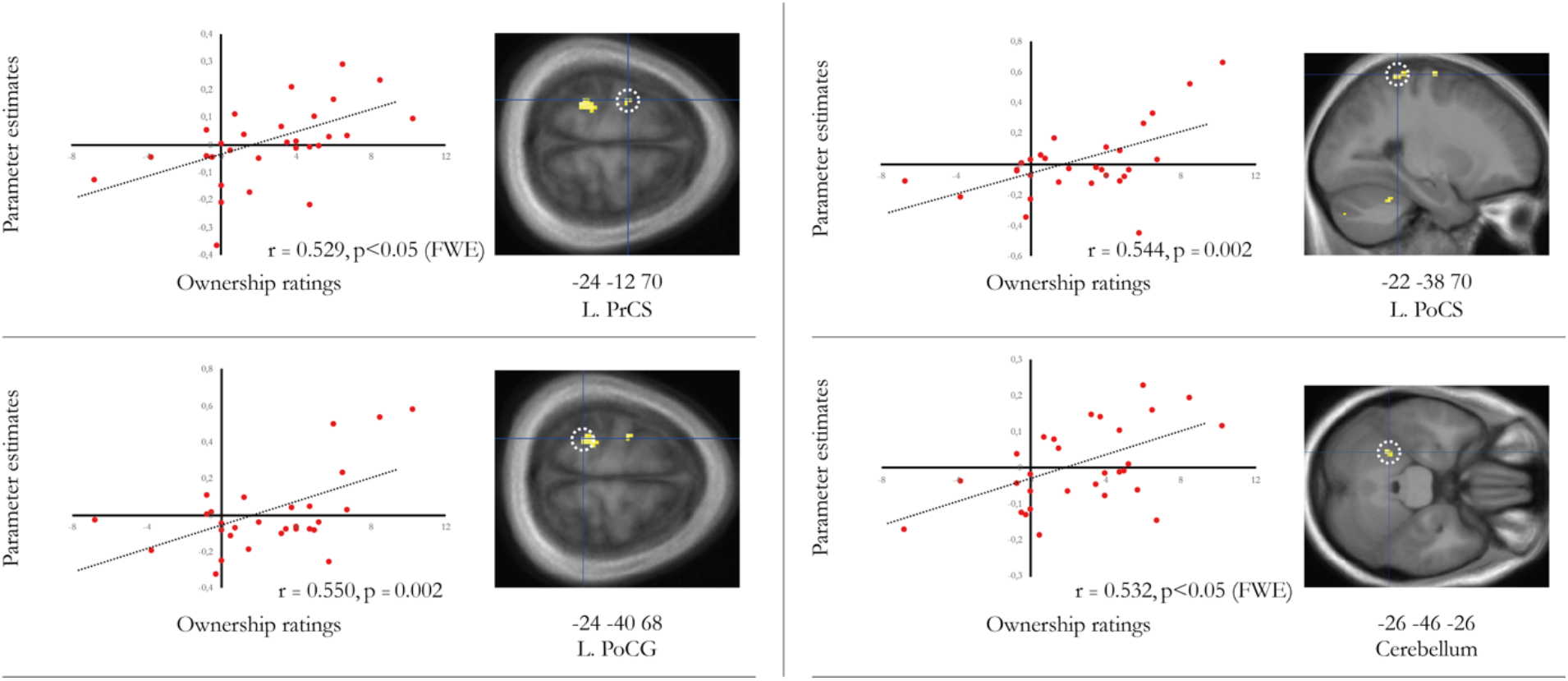
Correlation between behavioral ownership ratings (x-axis) and parameter estimates (y-axis, a.u.) in the left precentral sulcus (PrCS, -24, -12, 70), left postcentral gyrus (PoCG, -24, -40, 68), left postcentral sulcus (PoCS, -22, -38, 70) and left cerebellum (−26, -46, -26). Pearson’s r and p-values are given in each respective correlation plot. The peaks are displayed as activation maps (p<0.005, uncorrected) on representative sections of an average anatomical section and indicated with a dotted white line.

### The sense of agency is associated with activity in the left precentral and postcentral gyrus as well as right superior temporal gyrus

We then moved on to examine activations that reflect the sense of agency, that is, increases in activity dependent on actively generated movements as well as synchronous sensory feedback from the moving limb irrespectively if the hand was experienced as part of one’s body or not. To this end, we used the contrast [(A_M_S_T_C_O_–P_M_S_T_C_O_)–(A_M_A_T_C_O_–P_M_A_T_C_O_)] + [(A_M_S_T_I_O_–P_M_S_T_I_O_)–(A_M_A_T_I_O_–P_M_A_T_I_O_)] that represents agency across the congruent and incongruent conditions. In line with our hypotheses, we observed a significant activation peak in the left premotor cortex (−38, -8, 62; p<0.05, FWE corrected; Fig. 7; Table 2) and an activation in the right superior temporal gyrus that almost reached significance (58, -24, 12; p=0.051, FWE corrected; Fig. 7; Table 2). We also noted increases in activity in the intraparietal cortex bilaterally as well as the left superior temporal gyrus and left post central gyrus (p<0.001, uncorrected), but these activations did not survive correction for multiple comparisons and are only mentioned for purely descriptive purposes.

**Figure 7.**
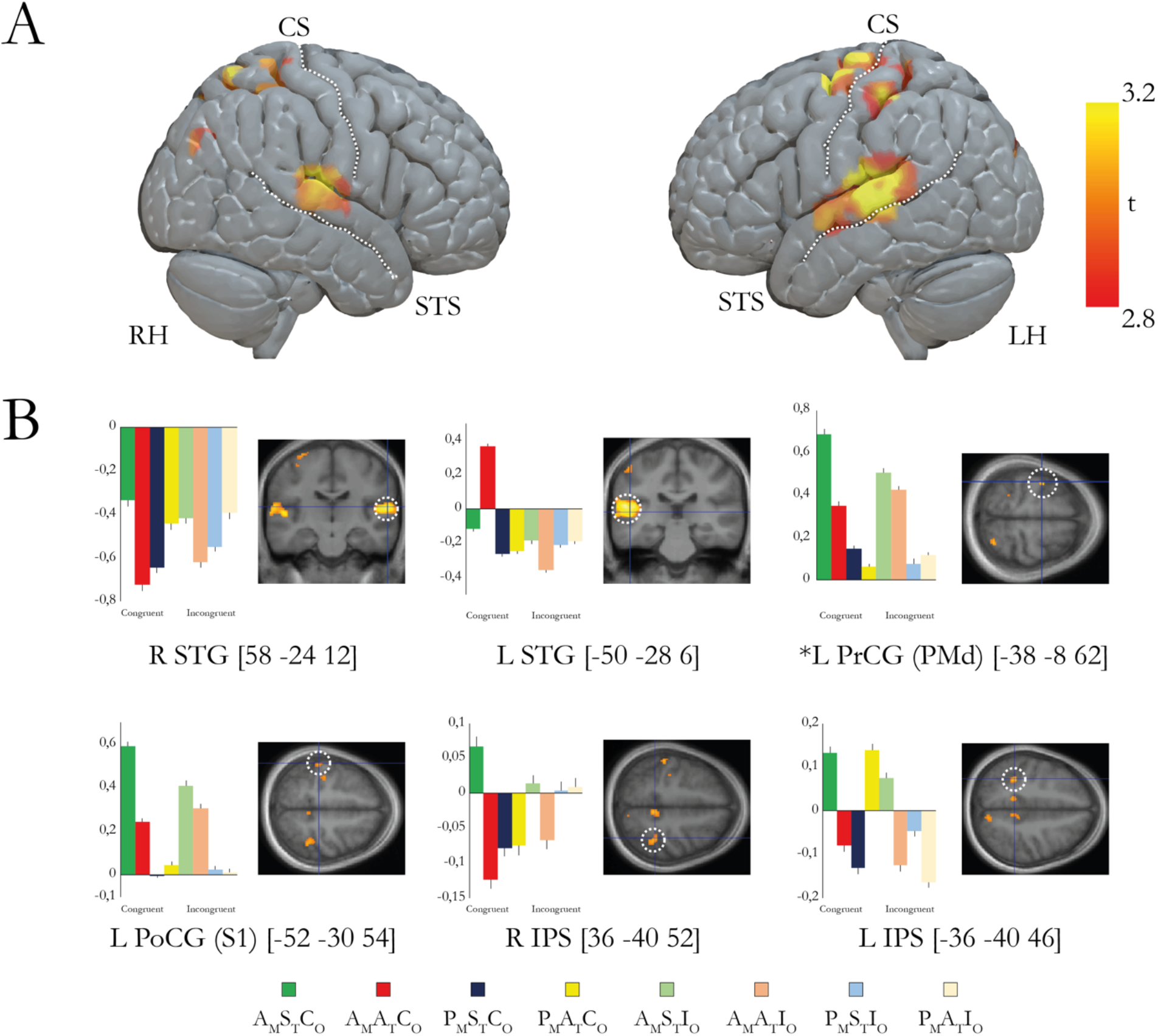
**A**. Overview of the brain regions that display activation reflecting the sense of agency. For display purposes only, the activations are projected on to a three-dimensional render of a standard brain with at a threshold of p<0.005 (uncorrected for multiple comparisons, k≥ 5). RH/LH, right/left hemisphere. STS, superior temporal sulcus. CS, central sulcus. **B**. Bar charts displaying the parameter estimates (a.u.) and SEs for the major peaks of activation. The coordinates are given in MNI space. The peaks are displayed in representative sections indicated by a dotted white circle on an activation map (p<0.005 uncorrected for display purposes). L/R, left/right. STG, superior temporal gyrus. PrCG, precentral gyrus. PoCG, postcentral gyrus. IPS, intraparietal sulcus. * indicates activation peaks that survive small volume correction (p<0.05 corrected); the peaks without an astrerix did not survive small volume correction and are reported in table 2 with their uncorrected p-value.

#### Conjunction analysis: agency and ownership overlap in the precentral gyrus (PMd)

To test for areas that showed increases in activity reflecting both ownership and agency, we used a conjunction analysis with the two two-way interaction contrasts described above for ownership and agency (Friston et al., 1999) (Fig. 8, Panel A). The analysis revealed a significant activation peak in the precentral gyrus (PMd, -38, -8 62, p<0.05 FWE corrected; Fig. 8, Panel A).

**Figure 8.**
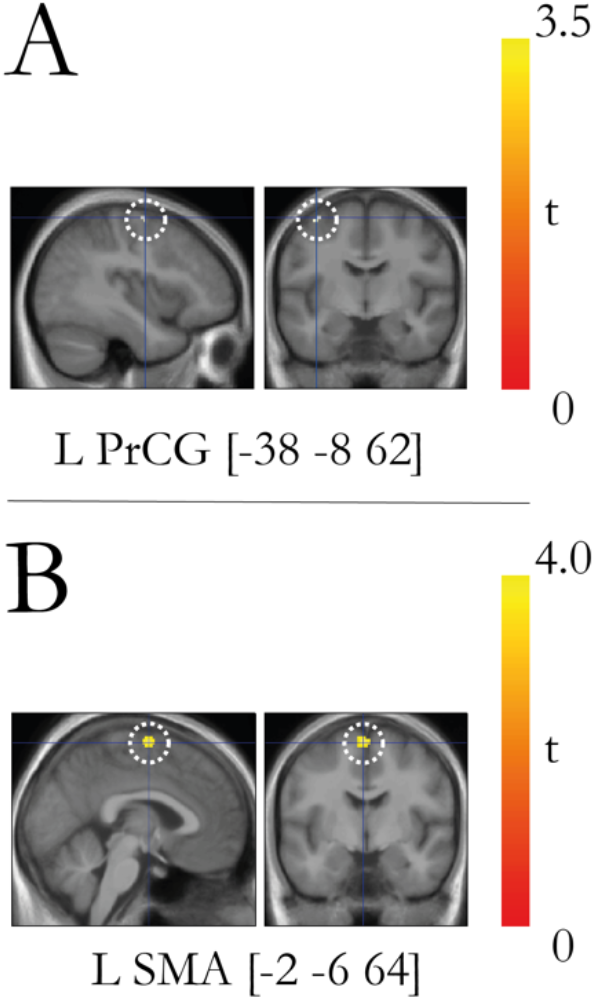
**A**. Conjunction analysis between the agency contrast and ownership contrast revealed overlapping activation in the left precentral gyrus (PMd). The peaks are displayed as activation maps (p<0.005, uncorrected) on representative sections of an average anatomical section and indicated with a dotted white line. **B**. PPI analysis of regions displaying increased connectivity with the seed region in the left postcentral gyrus (−38 -28 52). The left supplementary motor area (SMA) displays a task-specific increase in connectivity with the left post-central gyrus (S1). The peaks are displayed as activation maps (p<0.005, uncorrected) on representative sections of an average anatomical section and indicated with a dotted white line.

#### Interaction between ownership and agency revealed activation in somatosensory cortex

To test for interaction between ownership and agency we used the contrast [(A_M_S_T_C_O_–P_M_S_T_C_O_)– (A_M_A_T_C_O_–P_M_A_T_C_O_)]–[(A_M_S_T_I_O_–P_M_S_T_I_O_)–(A_M_A_T_I_O_–P_M_A_T_I_O_)]. This corresponds to the three-way interaction between movement type (active/passive), timing (synchronous/asynchronous) and rubber hand orientation (congruent/incongruent) and thus reveal neural responses unique to the combination of ownership and agency in the moving rubber hand illusion condition (A_M_S_T_C_O_). The results show significant activation in the left primary sensorimotor cortex with a significant peak of activation located in the postcentral gyrus at the level of the hand representations (−38, -28, 52; p<0.05, FWE corrected; Fig. 9) corresponding to the likely boundary of areas 1 and 3b; and three further peaks in the postcentral gyrus that did not survive corrections for multiple comparisons (p<0.005) (one possibly in area 3a close to the depth of central sulcus, p<0.005 uncorrected) (Fig. 9; Table 2).

**Figure 9.**
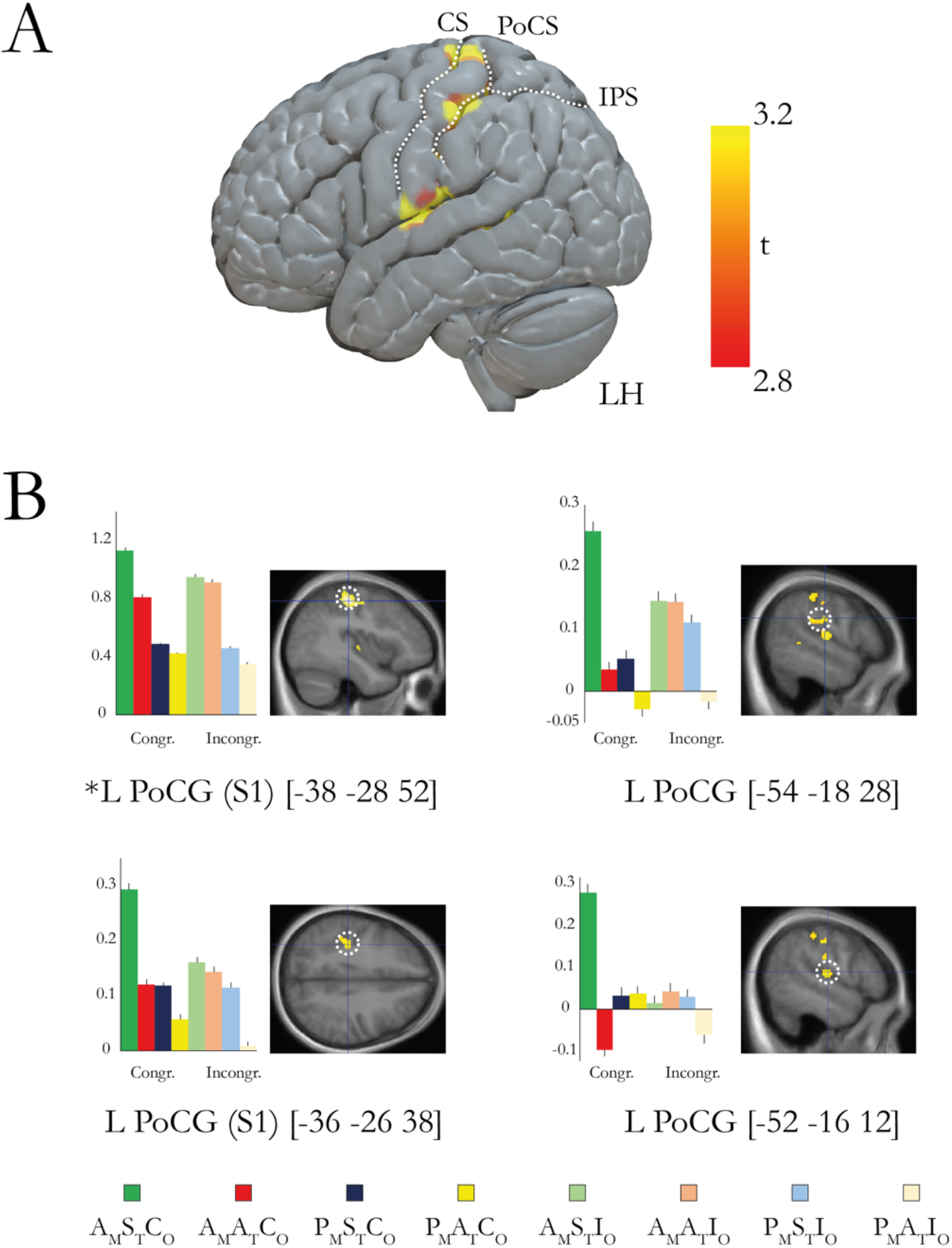
**A**. Overview of the brain regions that display activation reflecting increased activation related to agency of bodily objects compared to external objects. For display purposes only, the activations are projected on to a three-dimensional render of a standard brain with at a threshold of p<0.005 (uncorrected for multiple comparisons, k≥ 5). RH/LH, right/left hemisphere. IPS, intraparietal sulcus. PoCS, post-central sulcus. CS, central sulcus **B**. Bar charts displaying the parameter estimates (a.u.) and SEs for the major peaks of activation. The coordinates are given in MNI space. The peaks are displayed in representative sections indicated by a dotted white circle on an activation map (p<0.005 uncorrected for display purposes). L/R, left/right. PoCG, post-central gyrus. * indicates activation peaks that survive small volume correction (p<0.05 corrected); the peaks without an astrerix did not survive small volume correction and are reported in table 2 with their uncorrected p-value.

We should clarify here that the somatosensory activation under discussion can probably not be explained by somatosensory attenuation (Kilteni & Ehrsson, 2017, 2020; Zeller et al., 2014) or gating (Angel & Malenka, 1982; Kilteni & Ehrsson, 2022; Post et al., 1994; Voudouris et al., 2019), because we observed an increase in activity, not a reduction. Moreover, we also controlled the frequency and amplitude of the movements (Supplementary table 3 for source data), so it is unlikely that low-level differences in motor output or somatosensory feedback confounded our S1 findings in the three-way interactions because such factors were matched and thus eliminated in the statistical contrast.

In a post-hoc descriptive analysis of the effective connectivity in the three-way interaction of the factors timing, movement type and orientation congruency, we investigated the task-specific connectivity between the post-central gyrus, in a region corresponding to the primary sensory area (−38 -28 52) and the rest of the brain. We found that the sense of ownership in the presence of a sense of agency (or a sense of agency in the presence of a sense of ownership, depending on how one interprets the three-way interaction) increased the functional coupling between the left primary sensory cortex and the ipsilateral supplementary motor area (SMA) (Fig. 8, Panel B)

Next, we examined the opposite direction of three-way interaction contrast of movement type, movement type, synchrony, and orientation [(A_M_S_T_C_O_–P_M_S_T_C_O_)–(A_M_A_T_C_O_–P_M_A_T_C_O_)]–[(A_M_S_T_I_O_–P_M_S_T_I_O_)– (A_M_A_T_I_O_–P_M_A_T_I_O_)]. This contrast only revealed one activation in the left middle occipital gyrus and one smaller activation in the right middle occipital gyrus (Fig. 10; Table 2), but none of which survived correction for multiple corrections.

**Figure 10.**
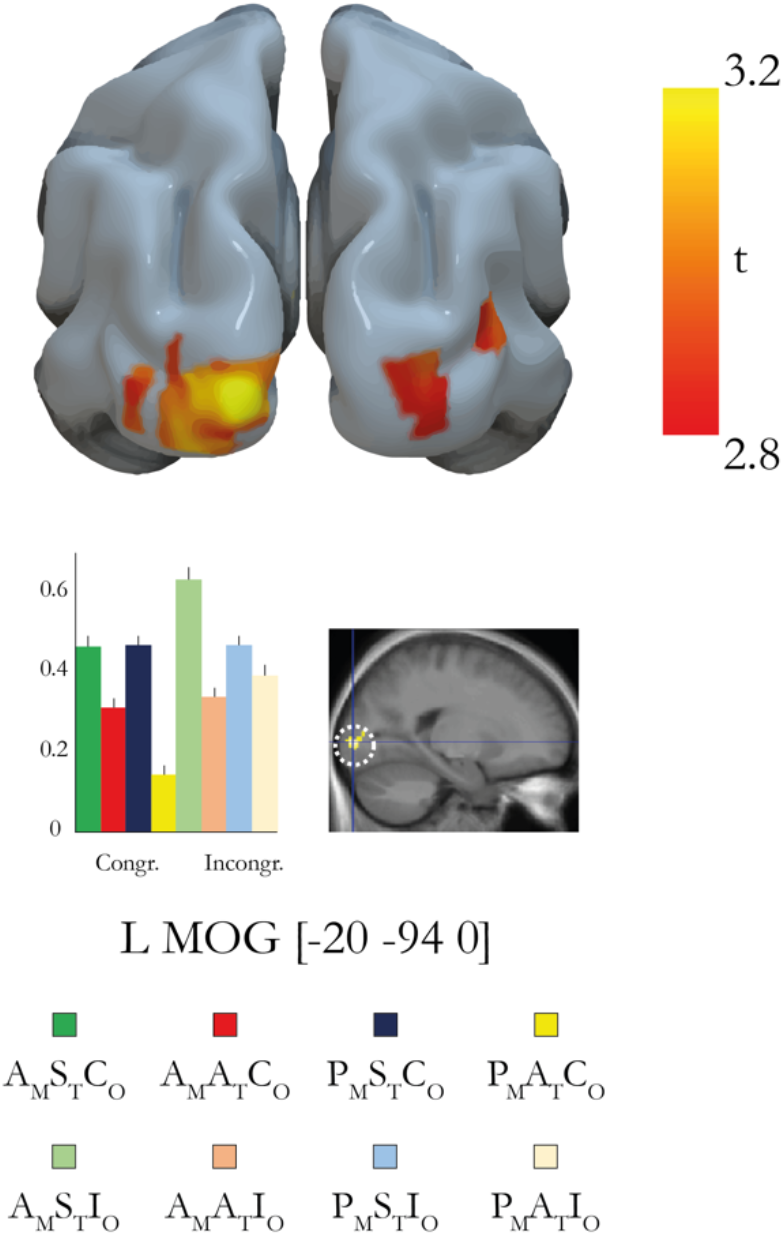
To investigate which brain regions are associated with the sense of agency of external objects as opposed to bodily objects, we defined a contrast that was the inverse of the three-way interaction [(A_M_S_T_C_O_– P_M_S_T_C_O_)–(A_M_A_T_C_O_–P_M_A_T_C_O_)]–[(A_M_S_T_I_O_–P_M_S_T_I_O_)–(A_M_A_T_I_O_–P_M_A_T_I_O_)]. The results show activation in the left middle occipital gyrus (p<0.001 uncorrected that does not survive correction for multiple comparisons) and right middle occipital gyrus (p=0.002, uncorrected). The coordinates are given in MNI space. L/R, left/right. MOG, middle occipital gyrus. The peak is displayed in a representative section and indicated by a dotted white circle on an activation map (p<0.005 uncorrected for display purposes, k≥ 5). The bar chart represents the parameter estimates (a.u.) for the peak.

#### Activations in insular cortex and right temporoparietal cortex reflect visuo-proprioceptive synchrony and asynchrony, respectively

In the previous literature it has been suggested that the right angular gyrus located in the temporoparietal region is involved in the loss of agency when there is a mismatch between the expected sensory consequences of self-generated movement and the sensory feedback (Farrer et al., 2003; Farrer & Frith, 2002; Tsakiris et al., 2010). Furthermore, it has been reported that the insular cortex show increases in activation when people experience agency (Farrer et al., 2003; Farrer & Frith, 2002). However, in our main planned contrasts reported above we did not find any changes in activation in these two regions, even at the level of uncorrected p-values (p<0.005). To examine this apparent inconsistency a bit further, we looked at the main effect of synchrony [(A_M_S_T_C_O_+A_M_S_T_I_O_+P_M_S_T_C_O_+P_M_S_T_I_O_) – (A_M_A_T_C_O_+A_M_A_T_I_O_+P_M_A_T_C_O_+P_M_A_T_I_O_)] and main effect of asynchrony contrasts [(A_M_A_T_C_O_+P_M_A_T_C_O_+A_M_A_T_I_O_+P_M_A_T_I_O_) – (A_M_S_T_C_O_+P_M_S_T_C_O_+P_M_S_T_I_O_+A_M_S_T_I_O_)], i.e., areas that show greater activation when visual feedback and finger movements are synchronous or asynchronous irrespectively of the senses of ownership or agency (i.e. across active and passive movements and across anatomically congruent or incongruent hand orientation). Interestingly, we found a large and significant activation (t=3.66, p=0.022, FWE-corrected) located in the right angular gyrus of the TPJ-region (50, -50, 32) that reflected the asynchronous relation between movement and visual feedback (main effect of asynchrony; Fig. 11; panel A). In contrast, synchrony of finger movements and visual feedback of the model hand’s finger movement (main effect of synchrony) was associated with significant activation (t= 3.71, p=0.020, FWE-corrected) of the left insular cortex (−38, -2, 10; Fig. 11; panel B). Thus, rather than reflecting the sense of agency or the loss of agency by mismatching sensory feedback, our results suggest that the insular cortex and right temporoparietal cortex are involved in a basic detection of synchronous or asynchronous multimodal stimuli.

**Figure 11.**
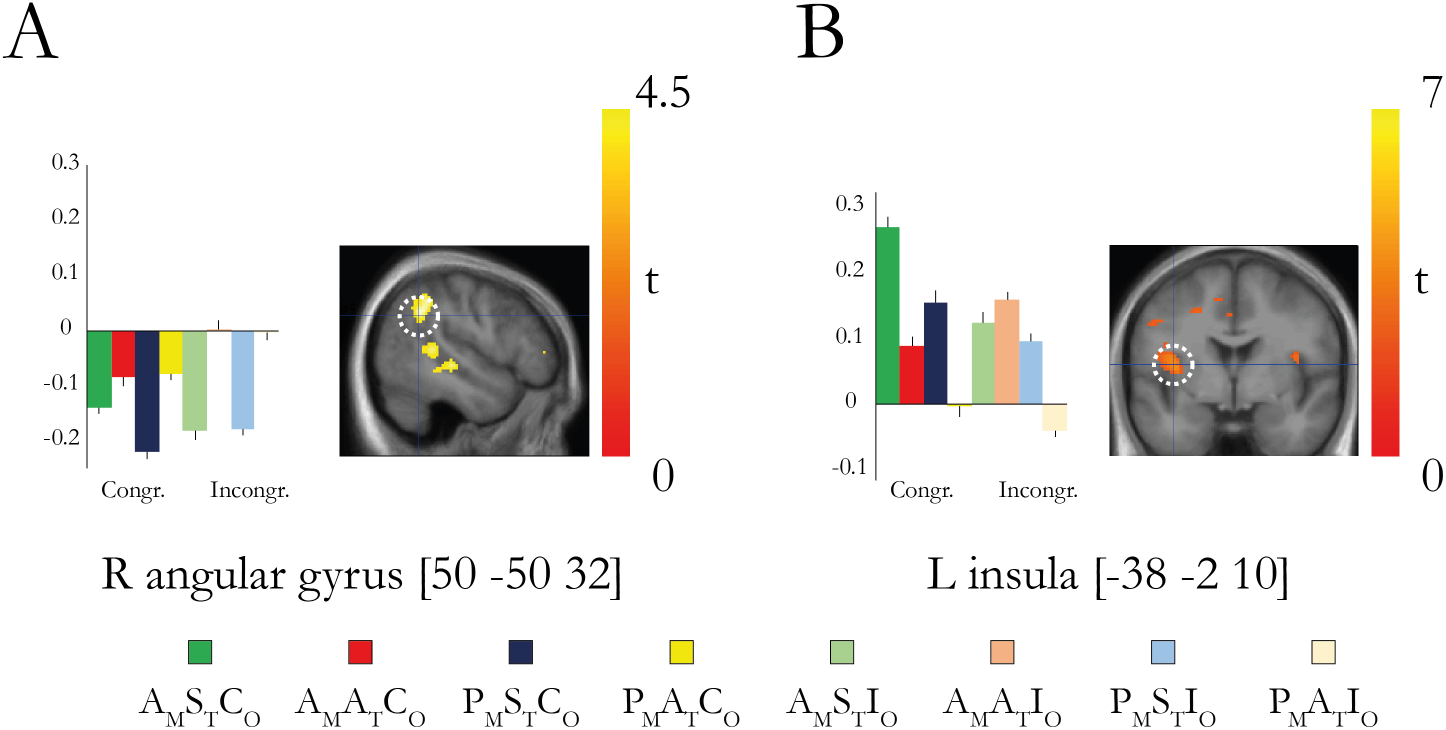
**A**. Activation in the right angular gyrus represented by the main effect of asynchrony (A_M_A_T_C_O_+P_M_A_T_C_O_+A_M_A_T_I_O_+P_M_A_T_I_O_) – (A_M_S_T_C_O_+P_M_S_T_C_O_+P_M_S_T_I_O_+A_M_S_T_I_O_) **B**. Activation in the left insular cortex represented by the main effect of synchrony (A_M_S_T_C_O_+P_M_S_T_C_O_+P_M_S_T_I_O_+A_M_S_T_I_O_) – (A_M_A_T_C_O_+P_M_A_T_C_O_+A_M_A_T_I_O_+P_M_A_T_I_O_). The coordinates are given in MNI space. The peak is displayed in a representative section and indicated by a dotted white circle on an activation map (p<0.005 uncorrected for display purposes).

#### Activations in supplementary motor cortex reflects main effect of active vs passive movements

Another area suggested to be involved in agency in previous fMRI studies, including agency in the moving RHI (Tsakiris et al 2010), is the supplementary motor area (SMA). However, this area did now show any agency-related activity in our agency contrast described above, not even at p<0.005 uncorrected. However, when we examined the main effect of movement type, contrasting all active versus all passive movement conditions in the current design, we observed significant activation of the SMA (A_M_S_T_C_O_+A_M_A_T_C_O_+A_M_S_T_I_O_+A_M_A_T_I_O_) - (P_M_S_T_C_O_+P_M_A_T_C_O_+P_M_S_T_I_O_+P_M_A_T_I_O_) (Fig. 12). Thus, this region seems to be important for generating movements voluntarily, thus implying it in movement planning, programming and volition more generally (Fried et al., 1991; Makoshi et al., 2011; Roland et al., 1980), but we found no evidence for specific involvement in sense of agency of the moving rubber hand.

**Figure 12.**
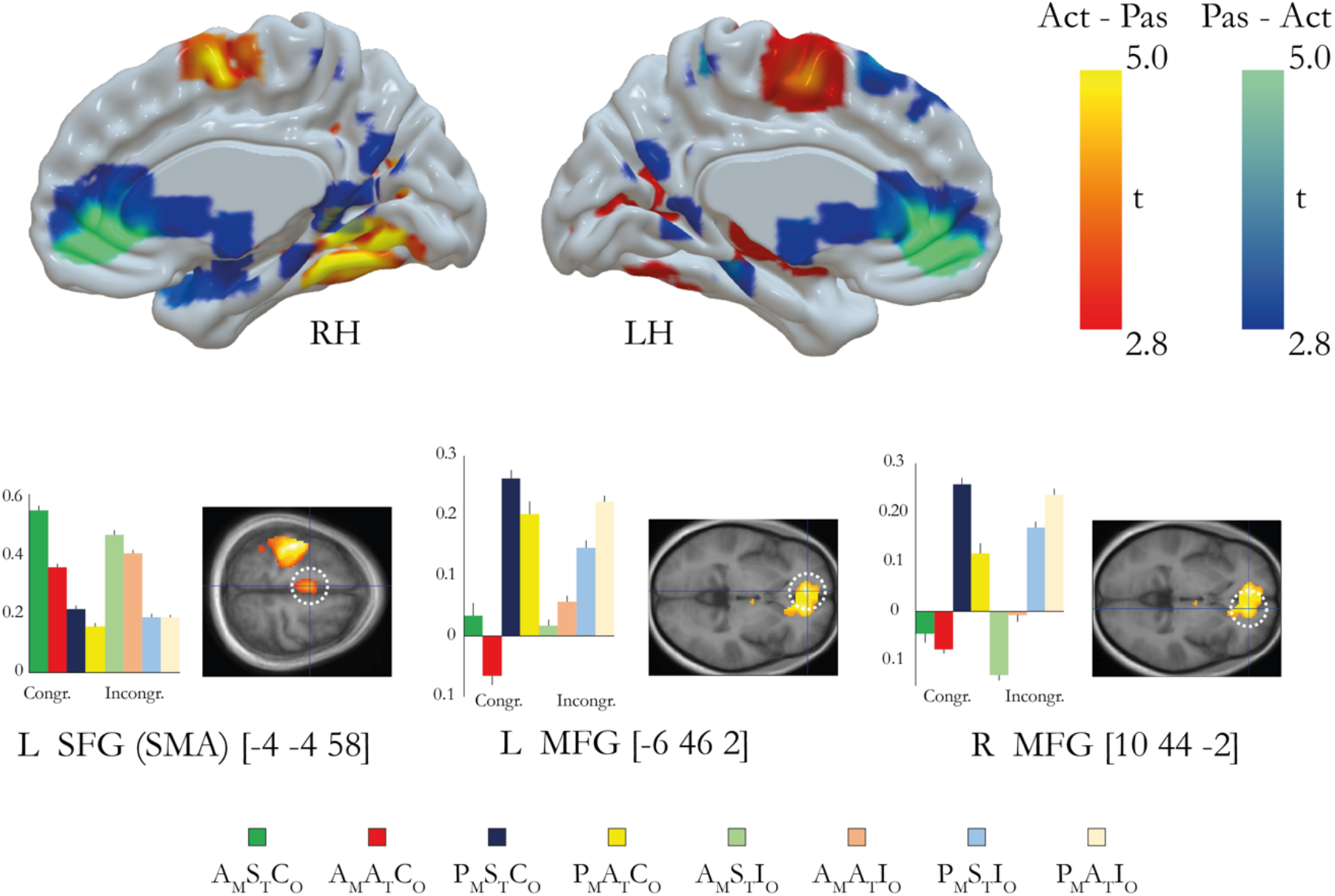
We defined a subtraction contrast (A_M_S_T_C_O_+A_M_A_T_C_O_+A_M_S_T_I_O_+A_M_A_T_I_O_) - (P_M_S_T_C_O_+P_M_A_T_C_O_+P_M_S_T_I_O_+P_M_A_T_I_O_) between all active condition and all passive conditions to serve as an internal control for the experimental setup and data acquisition. The contrast revealed activations in the left supplementary motor area (−4, -4, 58) as well as suprathreshold, significant activations in the left precentral gyrus (PMd; -42, -10, 60; t=7.82, p<0.001, FDR corrected, not shown), left precentral gyrus (M1; -40, -18, 56; t=9.20, p<0.011, FDR corrected, not shown) and right cerebellum (lobule VI; 20, -50, -24; t=9.23, p<0.001, FDR corrected, not shown) as well as left thalamus (−14, -22, 4; t=5.90, p=0.026, FDR corrected, not shown) and right angular gyrus (34, -50, 24; t=5.79, p=0.033, FDR corrected, not shown). Although we didn’t measure the EMG activity inside the MR-scanner, these results at least indicate that participants were more relaxed in the passive conditions than in the active conditions. We also ran the inverse contrast, (P_M_S_T_C_O_+P_M_A_T_C_O_+P_M_S_T_I_O_+P_M_A_T_I_O_)-(A_M_S_T_C_O_+A_M_A_T_C_O_+A_M_S_T_I_O_+A_M_A_T_I_O_) to identify regions that display an increase in neural activity associated with the passive conditions compared to the active conditions. Out results show activity in the bilateral medial frontal cortex. RH, right hemisphere. LH, left hemisphere. SFG= superior frontal gyrus, MFG= medial frontal gyrus.

When we looked for areas showing greater activity in the passive movement conditions than the active ones, we found a large activation in the medial prefrontal cortex in a region associated with default mode activity, autobiographical episodic memory, and self-related information processing (Tacikowski et al., 2017). The most straightforward interpretation is that, since participants did not have an active task in this condition (they just relaxed their hand and the experimenter generated the finger movements), the activity was higher in the default mode and therefore explaining the relatively higher activity in this medial prefrontal region compared to the active movement conditions when the participant had a task to move their finger repeatedly. This activation also corresponds well to similar activity observed in the passive finger movement condition in the study of Tsakiris et al 2010, which these authors attributed to ownership (Fig. 12).

#### Controlling for the number and frequency of taps in the different conditions

Using the optical sensor placed under the index finger of the participants, the number of taps as well as frequency of taps for each condition could be analyzed. The analysis was done on the time periods included in the fMRI analysis (i.e., excluding the time before illusion onset and corresponding time periods for conditions without illusion). The analysis revealed no significant differences across conditions in terms of frequency of taps (mean: 1.53 Hz; F=0.636, df=7, p=0.725), or the total number of taps (mean: 549; F=0.563, df=7, p=0.785).

## Discussion

This study has three main novel findings. First, the neural substrates of ownership and agency during movement were largely distinct, with body ownership associated with increases in activity in premotor cortex, posterior parietal and cerebellar regions, and the sense of agency related to increased activity in the superior temporal cortex and dorsal premotor cortex. Second, one active section of the dorsal premotor cortex was associated with both agency and body ownership, indicating a cortical site where ownership and agency information may be combined. Third, there was an interaction between body ownership and agency in the somatosensory cortex so that its activity was higher when participants experienced both sensations. This was accompanied by higher ownership ratings suggesting an agency-induced ownership enhancement of somatosensory cortical activity specific for voluntary movement. Collectively, these findings extend our knowledge of the neural basis of body ownership and agency and reveal their functional interaction and the relative neuroanatomical overlap and segregation during simple movement, which advances our understanding of how bodily self-consciousness is implemented in the human brain.

### The sense of body ownership during movement: integration of spatio-temporally congruent visuo-proprioceptive signals in premotor-parietal-cerebellar regions

The present study extends the previous neuroimaging literature on the neural basis of body ownership (Brozzoli et al., 2012; H. H. Ehrsson, 2007; H. H. Ehrsson et al., 2004; Gentile et al., 2013; Guterstam, Collins, et al., 2019; Guterstam et al., 2013; Limanowski & Blankenburg, 2016; Petkova et al., 2011; Preston & Ehrsson, 2016) into such experience arising from the sensory feedback of movement. The sense of ownership of the moving rubber hand was associated with significant activations in the left premotor cortex (precentral gyrus), posterior parietal cortex (left supramarginal gyrus) and the right lateral cerebellum. These activations probably reflect the integration of spatially and temporally congruent visual information from the moving rubber hand and kinesthetic-proprioceptive information from the hidden real hand because the neural response was specifically related to the conditions when the rubber hand was placed in an anatomically congruent condition and the seen and felt movements synchronous; controlling for agency effects and effects related to active versus passive movement.

The difference between visuo-kinesthetic integration, studied here, and visuo-tactile integration, investigated in previous RHI studies, can probably explain the differences in precise localization of the activation peaks in the premotor cortex compared to previous studies (e.g. (H. H. Ehrsson et al., 2004)). Although activations has been seen in both ventral and dorsal aspects of the premotor cortex in previous RHI studies (Gentile et al., 2013; Guterstam, Collins, et al., 2019), the most consistent activations tend to be have been located in the ventral premotor cortex (H. H. Ehrsson et al., 2004; Gentile et al., 2013; Grivaz et al., 2017; Guterstam, Collins, et al., 2019; Guterstam et al., 2013; Limanowski & Blankenburg, 2016). The dorsal premotor cortex is active during passive hand and arm movements, finger tapping (Bengtsson et al., 2009; Ullén et al., 2003), and illusory hand and arm movements triggered by muscle tendon vibration (Naito et al., 1999, 2005), consistent with a role in multisensory representation of the upper limb in space. The current activation in the supramarginal gyrus (p<0.05 corrected) is consistent with earlier body ownership illusion studies based visoutactile stimulation (Gentile et al., 2011, 2013; Petkova et al., 2011), and the current intraparietal cortex activation is located in a section of this sulcus associated with multisensory integration in perihand space (Brozzoli et al., 2011; Lloyd et al., 2003; Makin et al., 2007). We also observed activity in the ipsilateral lateral cerebellum in line with previous fMRI studies on various versions the rubber hand illusion based on visuotactile stimulation (H. H. Ehrsson et al., 2004, 2005; Guterstam et al., 2013) and limb-movement illusions (H. H. Ehrsson et al., 2005; Hagura et al., 2009). Importantly, the current findings extend the previous literature on body ownership and body representation by demonstrating a role for these premotor-parietal-cerebellar regions in the sense of limb ownership during movement.

### The sense of agency in own bodily movement: premotor and superior temporal cortex

We could isolate activity in the dorsal premotor cortex and superior temporal cortex reflecting agency over limb movement while controlling for unspecific effects related to multisensory synchrony-asynchrony detection, active versus passive movement, and body ownership. The dorsal premotor area has been reported in previous studies on the sense of agency over sensory events caused by voluntary movement (David et al., 2008; Haggard, 2017; Nahab et al., 2011; Sperduti et al., 2011; Yomogida et al., 2010) so our finding extends this to agency over perceived own bodily movement. The dorsal premotor cortex is anatomically connected to, and receives input from, the dorsolateral prefrontal cortex regarding intentions and the initiation of voluntary action in the context of an overall action plan (Abe & Hanakawa, 2009; Koechlin et al., 2003; Passingham, 1993; Yamagata et al., 2012) and receives multisensory input from the posterior parietal cortex regarding one’s own body as well as external sensory events; the dorsal premotor area can also influence movement execution in M1 and receive feedback from this area through direct cortico-cortical connections (Dum et al., 2002; Porter & Lemon, 1995). The dorsal premotor cortex is thus in an excellent position, anatomically and physiologically, to play a central role in the sense of agency by integrating and comparing signals related to voluntary motor commands and sensory feedback, consistent with our findings.

Interestingly, the section of the dorsal premotor cortex associated with agency also showed body ownership-related activity, as revealed in our conjunction analysis. This finding suggest that the neural basis of body ownership and agency are not completely distinct (Tsakiris et al., 2010), and there exists at least one cortical area involved in both processes. Different neuronal populations within the dorsal premotor cortex could implement the formation of a coherent multisensory representation of the hand in space (ownership) and generation of voluntary motor commands and the matching of those commands’ outcomes with the sensory feedback and predictions (agency), or the same neuronal population within this area may implement both these mechanisms (which could be tested in future studies with BOLD adaptation or multivoxel pattern analysis). Regardless of which, our findings suggest a more intimate relationship of the representations of body ownership and agency in the premotor cortex than commonly assumed and indicate that more attention should be given to this region in future studies on the neural mechanisms of agency of bodily action.

Previous neuroimaging studies have implied the superior temporal cortex in the sense of agency, but reported activation in superior temporal gyrus reflecting the *loss* of agency when controlling a virtual limb (Nahab et al., 2011; Uhlmann et al., 2020). However, these studies did not control for multisensory synchrony-asynchrony, the visual appearance (and identity) of the hand, or body ownership. In contrast, we found a relative activity *increase* that reflected *gaining* agency of the moving rubber hand, although all experimental conditions were deactivated compared to the resting baseline. The current activation peak is located more ventral and anterior to the deactivations in the previous studies (Nahab et al., 2011; Uhlmann et al., 2020), making direct comparisons difficult. Although, the precise functional role of the superior temporal cortex in agency is unclear, this region has been associated with action observation (Kilintari et al., 2014), visual processing of biological motion (Saygin, 2007), and perception of causality between sensory events (Blakemore et al., 2001) which collectivity points towards a function of supporting the (visual) perception of causality relationships between the seen finger movement and the executed finger action, which presumably is an important component of the agency experience.

### Interaction of body ownership and agency in somatosensory cortex

Our analysis revealed somatosensory activity that was uniquely related to the situation when both ownership and agency were experienced over the moving rubber hand (interaction effect). In principle, this activity could reflect a change in body ownership caused by agency or a change in agency caused by ownership. We think the former is more likely because the behavioral data showed a corresponding interaction effect in the questionnaire hand-ownership ratings (no such interaction was found in the agency ratings). Thus, the somatosensory activity may be related to a change in the somatic feeling of the rubber hand illusion when this illusion is produced by visuo-motor correlations during active finger tapping as opposed to passive finger movements and visuo-kinesthetic correlations. Motor commands and efferent signals can influence limb movement sensations (Gandevia et al., 2006; Walsh et al., 2010), and thus we theorize that efferent information related to the active voluntary motor commands made the ownership experience more vivid by boosting kinesthetic sensations from the rubber hand’s finger movements. Such efferent signals could originate from premotor areas and influence the somatosensory cortex via cortico-cortical connections, which is supported by the finding of increased effective connectivity between the SMA and S1 during the three-way interaction (Fig. 8). Alternatively, the somatosensory activity changes could stem from cerebellar influences expressed via cortico-cerebellar connections (Hagura et al., 2009; Kilteni & Ehrsson, 2020), perhaps reflecting somatosensory predictions specific for own finger movements. As said, the somatosensory activity might also reflect a special component of agency over one’s bodily movements – “bodily agency” – perhaps reflecting differences between self-related somatosensory predictions and predictions about external (e.g., visual) events that are indirectly caused by voluntary action. Regardless of the underlying mechanism and conceptualization as changes in body ownership or (bodily) agency, our finding links somatosensory activity to the combination of ownership and agency during voluntary limb movement.

## Acknowledgements

Zakaryah Abdulkarim was funded by a PhD student grant from the Karolinska Institutet (Clinical Scientist Training Programme). Henrik Ehrsson and project costs were supported by funding from the Swedish Research Council, The Göran Gustafsson Foundation, the Torsten Söderbergs Stiftelse and Hjärnfonden. Arvid Guterstam was supported by the Wenner-Gren Foundations and the Swedish Brain Foundation. We thank Martti Mercurio for help with the experimental setups and the staff at the Karolinska University Hospital Solna’s MR-center for support with the fMRI. We thank Konstantina Kilteni for valuable input during the analysis of the fMRI data.

## Notes

### Competing Interest Statement

The authors have declared no competing interest.

